# CD38 defines a therapeutically targetable pathogenic T cell population for precision immunotherapy in autoimmune diabetes

**DOI:** 10.64898/2026.06.27.734105

**Authors:** Shiva Pathak, Rizwan Ahmed, Nadine Nagy, Sooyeon Lee, Cameron S. Bader, Shobha Regmi, Bettina P. Iliopoulou, Pin-I Chen, Biki Gupta, Alejandro Villar-Prados, Yong Bin Kim, Noor Hussein, Emily SooHoo, Abigail Twoy, Avnesh S. Thakor, Kent P. Jensen, Paul J. Utz, Mark M. Davis, Justin P. Annes, Everett H. Meyer

## Abstract

Type 1 diabetes (T1D) is caused by T cell-mediated autoimmune destruction of insulin-producing islet β-cells. Treatment with T-cell depleting therapies delays the progression of stage 2 and 3 T1D, but these agents exert broad immunosuppressive effects on T cell populations, including T regulatory cells (Tregs), which are key in promoting immune tolerance. We evaluated non-obese diabetic (NOD) mice and recently diagnosed T1D patients and identified CD38 as a marker for pathogenic T cell populations. Using adoptive T-cell transfer in Recombination Activating Gene 1 knockout NOD mice and in a humanized mouse model of autoimmune diabetes, we demonstrated that CD38-expressing autoreactive T cells drive diabetes pathogenesis. Furthermore, we found that selective depletion of CD38^+^ cells, using an anti-CD38 monoclonal antibody (mAb), prevents insulitis and diabetes onset without depleting CD4^+^CD25^+^ Tregs. Administration of anti-CD38 mAb did not adversely affect islet function and may selectively eliminate immunogenic senescent islet β-cells. These results support the strategy of selectively depleting diabetogenic T cells using an anti-CD38 mAb to treat T1D and restore immune tolerance. Therefore, transient depletion of autoreactive T cells using anti-CD38 mAb may provide a novel strategy to prevent or abrogate β-cell autoimmunity in T1D.

**One sentence summary:** Pathogenic autoreactive are characterized by CD38 overexpression, and their selective depletion with anti-CD38 monoclonal antibody prevents autoimmune diabetes.

## INTRODUCTION

Type 1 diabetes (T1D) is an autoimmune disorder primarily caused by islet-infiltrating autoreactive T cells^1^. Efforts to prevent T1D have focused on broad T cell depletion that can modify disease course, but rarely yields a durable response^2,3^. Teplizumab, an anti-CD3 monoclonal antibody, is currently the only FDA approved therapy for early stage T1D and can delay disease onset in at-risk individuals^4^. Despite the clinical use of Teplizumab, patient response is limited, heterogenous, and not durable^5,6^. Beyond teplizumab, broader T cell depletion with anti-thymocyte globulin (ATG) and B cell depletion with rituximab (anti-CD20) have failed to yield durable responses^7–9^. In addition, these lymphodepletion strategies have broad immunosuppressive effects that adversely affect regulatory T cells (Tregs) and other immune effector populations^10,11^. Thus, it is necessary to develop better targeted therapeutic strategies that promote durable immune tolerance and overall better clinical outcomes in T1D.

CD38 is a multifunctional type II transmembrane glycoprotein that acts as an ectoenzyme with nicotinamide adenine dinucleotide (NAD+) hydrolase and adenine diphosphate (ADP)-ribosyl cyclase activities, generating secondary messengers such as ADP ribose and cyclic-ADP ribose that regulate intracellular calcium mobilization^12^. In autoimmune and inflammatory diseases, elevated CD38 has been reported on activated T and B cells, plasmablasts, monocytes, and tissue-resident myeloid cells^13^. CD38 is upregulated by T cells during their activation, indicating increased metabolic and signaling capacity and potentiation of effector responses^14^. While upregulation of CD38 has been documented in multiple inflammatory contexts^13^, it is yet to be determined if pathogenic autoreactive T cells preferentially express CD38 compared to other immune cells. Furthermore, the functional significance of CD38 in autoreactive T cells has not been determined. Thus, elucidating the role of CD38 in pathogenic autoreactive T cells in T1D through integrated phenotypic and functional analyses is essential for determining whether CD38^+^ T cells drive autoreactive pathogenesis and to guide the development of novel targeted immunosuppressive therapies that spare physiologic immune populations, particularly Tregs.

In this study, we show the role of CD38^+^ autoreactive T cells in driving T1D pathogenesis and provide a rationale for targeting CD38 to selectively deplete pathogenic autoreactive T cells in T1D. We identified CD38 as a marker of pathogenic autoreactive T cells *via* bulk RNA sequencing and flow cytometric analysis of islet antigen-specific CD8^+^ T cells from diabetes-susceptible, female non-obese diabetic (NOD) mice. We also found that a short-course anti-CD38 monoclonal antibody (mAb) prevents insulitis and disease progression in NOD mice. Furthermore, we identified CD38^+^ autoreactive T cells among the recently diagnosed stage 3 T1D patients and the frequency of these cells decrease in blood of stage 4 T1D patients. In a humanized mouse model of T1D, targeting CD38^+^ T cells with a human anti-CD38 mAb mitigates insulitis. Overall, these data suggest that CD38^+^ autoreactive T cells are mechanistic drivers of diabetogenesis in T1D. Transient lymphodepletion using anti-CD38 mAb represent a novel therapeutic target for future clinical trials for stage 2 and 3 T1D patients and for patients with established T1D receiving islet transplant or other β-cell replacement therapies.

## RESULTS

### Phenotyping of autoreactive T cells reveals upregulated expression of CD38

To define therapeutic targets for the lymphodepletion of autoreactive T cells, we sort-purified islet-specific glucose-6-phosphatase catalytic subunit-related protein (IGRP, sequence: 206-214)-reactive CD8^+^ T cells from pooled spleens and draining lymph nodes of prediabetic NOD mice using IGRP H-2^kd^ tetramers and performed bulk RNA sequencing (**Fig 1A**). Transcriptomics were generated using bulk CD8^+^ T cells from age- and sex-matched NOD mice for comparison. The gating hierarchies for sorting IGRP-reactive CD8^+^ T cells are shown in **Fig 1B and Fig S1**. Principal component (PC) analysis revealed that IGRP-reactive CD8^+^ T cells formed a transcriptional cluster with higher intra-group consistency, whereas their profiles remained distinct from bulk CD8^+^ T cells (**Fig S2A**). Differential expression analysis identified 1304 upregulated and 413 downregulated genes among IGRP-reactive CD8^+^ T cells *versus* bulk CD8*^+^* T cells (**Fig S2B**). Hierarchical clustering and heatmap analysis highlighted proinflammatory and effector genes, including *Ccl5*, *Mki67*, *Klre1*, *Il18rap*, *Ifng*, *Cd38*, *Cxcr3*, *Ccl4*, *Ctla2a*, *Ccl3*, *Cx3cr1*, *Irf8*, *Cd101*, and *Ccr5* (**Fig 1C and Fig S2C**). Furthermore, gene set enrichment analysis demonstrated concurrent activation of multiple proinflammatory pathways, including those associated with complement, allograft rejection, IL-2 STAT5 signaling, IFN-α/IFN-γ responses, TNF signaling, and reactive oxygen species pathways; indicating a shift from a canonical CD8^+^ phenotype towards a proinflammatory and highly-activated status (**Fig 1D**). CD38 emerged as one of the top upregulated genes in autoreactive T cells compared to the bulk CD8^+^ T cells. Given the established role of CD38 as a marker and mediator of T cell activation in inflammatory and autoimmune settings, we prioritized CD38 as a candidate therapeutic target in autoreactive T cells. Elevated expression of CD38 in autoreactive T cells suggested an activated state that may sustain inflammatory signaling and promote the recruitment or activation of additional effector cells within the islet microenvironment, thereby contributing to β-cell destruction in T1D. To determine whether *Cd38* upregulation observed at the transcriptomic level was also evident at the protein level, we performed flow cytometric analysis of NOD splenocytes. Cells were stained with antibodies against CD3, CD8, CD25, and CD38. IGRP-reactive CD8^+^ T cells were identified using APC- and PE-labeled IGRP p206-214 peptide-loaded tetramers. Data confirmed upregulation of CD38 among IGRP-reactive CD8^+^ T cells. Representative flow cytometry contour plots (**Fig 1E**) and quantification of CD38 expressing cells showed a statistically significant (*p*<0.0001) increase in the percentage of CD38^+^ cells among the autoreactive T cells compared to bulk T cells or CD4^+^CD25^+^ Tregs (**Fig 1F**). These integrated transcriptomics and flow cytometric analyses identified a proinflammatory autoreactive CD8^+^ T cell population in prediabetic NOD mice that may promote early islet invasion and β-cell destruction.

**Fig 1.**
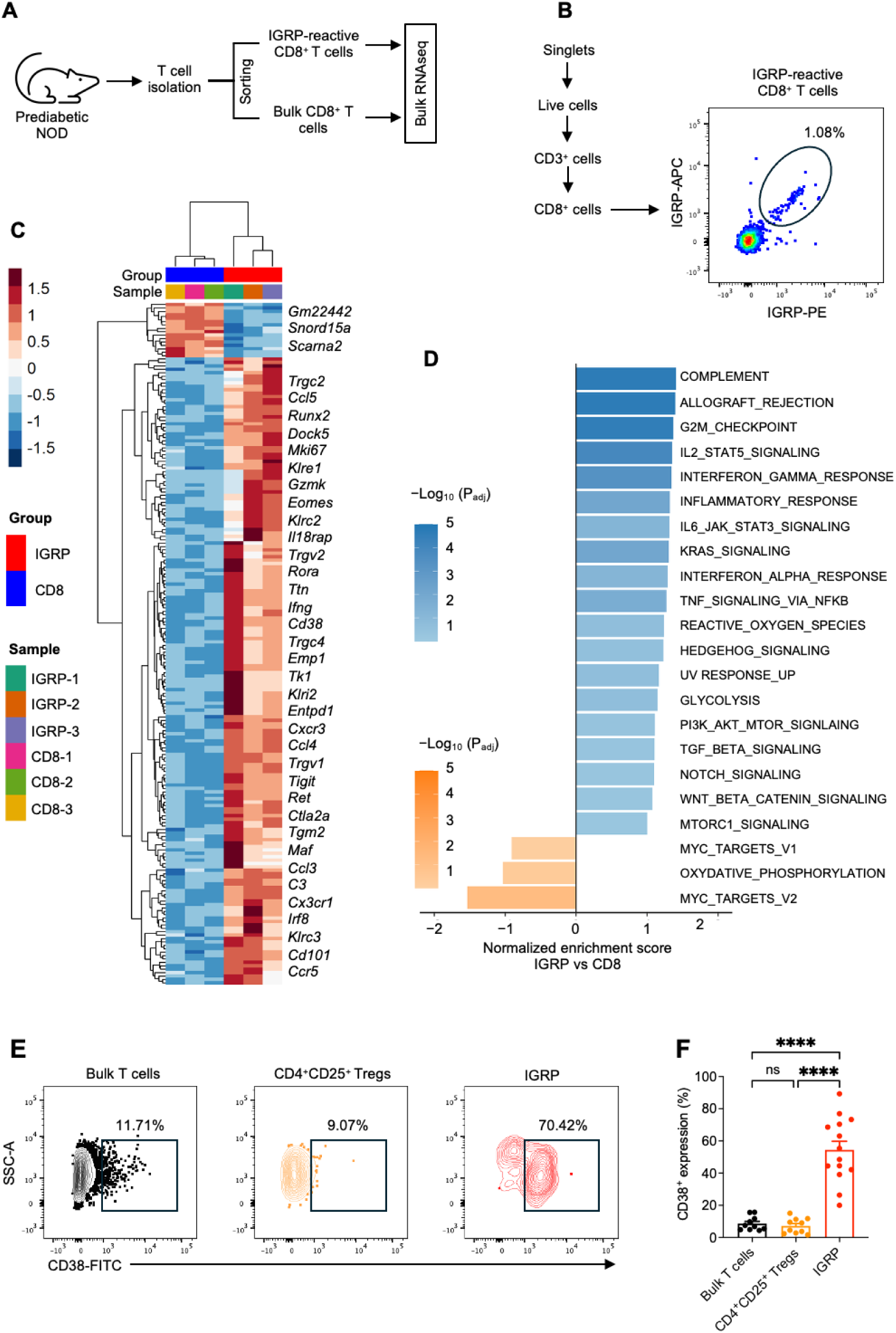
Autoreactive T cells in NOD mice have elevated CD38 expression. **(A)** IGRP-reactive CD8^+^ T cells were sorted from 10 weeks old prediabetic female NOD mice using IGRP H-2^Kd^ tetramers and processed for bulk RNA sequencing. IGRP-reactive CD8^+^ T cells and bulk CD8^+^ T cells in each sample were collected from spleens pooled from 5 NOD mice. (**B**) Gating hierarchy for sorting IGRP-reactive CD8^+^ T cells. (**C**) Heatmap and hierarchical clustering of top 200 most differentially expressed genes by adjusted *p*-values in IGRP *versus* bulk CD8^+^ T cells. (**D**) Gene-set enrichment analysis of IGRP *versus* bulk CD8^+^ T cells. The 20 most immune-related and differentially expressed pathways by normalized enrichment score are shown. (**E**) Representative flow cytometry contour plots from at least 3 independent experiments showing surface expression of CD38 in bulk T cells, CD4^+^CD25^+^ Tregs and IGRP-reactive CD8^+^ T cells. (**F**) Quantification of the percentage of CD38^+^ cells among bulk T cells, CD4^+^CD25^+^ Tregs, and IGRP-reactive CD8^+^ T cells. ns: not significant; \*\*\*\**p*<0.0001.

### Depletion of CD38^+^ T cells abrogates diabetes in adoptive transfer model using NRG mice

To test whether CD38^+^ autoreactive T cells drive islet autoimmunity in NOD mice, we conducted an adoptive T cell transfer experiment using 12 weeks old diabetes-susceptible NOD mice as T cell donors. We transferred NOD T cells or CD38-depleted NOD T cells (NOD-CD38^dep^) into Recombination Activating Gene 1 deficient (RAG1^−/−^)-NOD (NRG) recipients. We measured T cell persistence in the NRG recipient mice at day 20 post-T cell transfer. Non-fasting blood glucose (NBG) levels were monitored, and endpoint histological analysis was performed at day 60 post-transfer (**Fig 2A**). The frequency of total CD3^+^ T cells did not differ between the two groups on day 20 post-transfer (**Fig 2B**). The representative flow cytometry plots showing IGRP-reactive CD8^+^ T cells in NOD T cell and NOD-CD38^dep^ groups are depicted in **Fig 2C**. The frequency of IGRP-reactive CD8^+^ T cells was significantly lower in the NOD-CD38^dep^ group compared to NOD T cell group (*p*<0.05) (**Fig 2D**). Longitudinal NBG monitoring for up to 60 days revealed a striking difference in the disease incidence: 11/15 (73%) mice in the NOD T cell group became hyperglycemic, whereas only 1/15 (7%) mice in the NOD-CD38^dep^ group became hyperglycemic (**Fig 2E-F**). Kaplan-Meier analysis demonstrated a significantly higher (*p=0.0002*) diabetes-free survival in the NOD-CD38^dep^ group compared to the NOD T cell group (**Fig 2G**). To directly assess the extent of peri-islet immune infiltration, we collected pancreata at day 60 post-transfer and conducted hematoxylin and eosin (H&E) staining and immunostaining for T cells (CD3, CD38) and insulin. H&E staining of pancreata showed massive peri-islet lymphocytic infiltration in NOD T cell group, whereas NOD-CD38^dep^ group showed markedly reduced lymphocyte infiltration indicating decreased insulitis (**Fig 2H**). Further immunohistochemistry analyses confirmed that CD3^+^ T cells and CD38^+^ T cell infiltrates were substantially lower in the pancreata of the NOD-CD38^dep^ group compared to NOD T cell group (**Fig 2I-J**). Insulin staining demonstrated a loss of insulin-positive cells in the NOD T cell group, whereas insulin staining preserved in the NOD-CD38^dep^ group (**Fig 2K**). These data indicate that CD38^+^ autoreactive T cells constitute a pathogenic subset that drives islet infiltration and β-cell destruction, and that selective ablation of CD38^+^ T cells mitigates disease onset.

**Fig 2.**
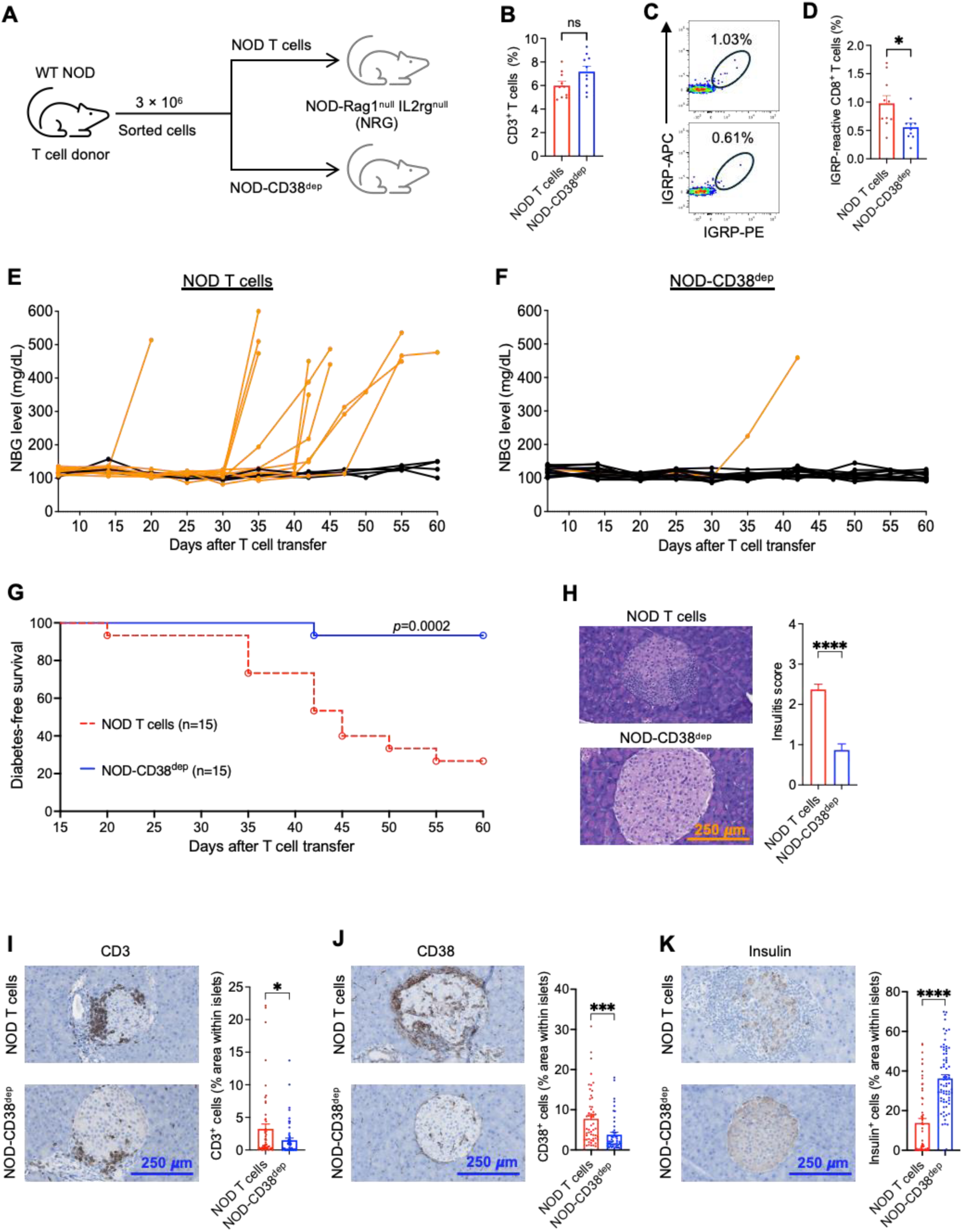
Depletion of CD38^+^ T cells prior to adoptive T cell transfer abrogates diabetes transfer in NRG mice. **(A)** CD3^+^ T cells were sorted from 12 weeks old prediabetic female NOD mice and transferred into NSG recipients. A total of 3 × 10^6^ T cells with or without CD38^+^ T cells were injected in each mouse. (**B**) Percentage of CD3^+^ T cells in recipient NRG mice at day 20 post-transfer. (**C**) Representative flow cytometry plots showing IGRP-reactive CD8^+^ T cells in recipient NRG mice at day 20 post-transfer (**D**) Percentage of IGRP-reactive CD8^+^ T cells in recipient NRG mice at day 20 post-transfer. (**E**) Non-fasting blood glucose (NBG) profiles of recipient NRG mice that received total CD3^+^ T cells. (**F**) NBG profiles of recipient NRG mice that received CD38-depleted CD3^+^ T cells (NOD-CD38^dep^). In **E** and **F**, each curve represents NBG profile of an individual mouse; orange curves indicate NBG profiles of mice that became hyperglycemic and black curves indicate NBG profiles of mice that remained euglycemic during the 60-day observation period. (**G**) Kaplan-Meier analysis showing diabetes-free survival of recipient NRG mice post-transfer. (**H**) Representative H&E staining of pancreata (left) and quantification of insulitis score (right) in recipient NRG mice at day 60 post-transfer. Insulitis score calculated from 35-40 islets in each group. 0: no insulitis, 1: peri-insulitis, 2: infiltrative insulitis <50% of the islet, 3: infiltrative insulitis >50% of the islet. (**I-K**) CD3, CD38, and insulin immunohistochemistry analysis in pancreas sections of recipient NRG mice at day 60 post-transfer. Data represent mean±SEM and were pooled from three independent experiments. Scale bar: 250 *µ*m; magnification: 40×. ns: not significant; \**p*<0.05; \*\*\**p*<0.001; \*\*\*\**p*<0.0001.

### Anti-CD38 mAb treatment prevents insulitis and hyperglycemia in NOD mice

To test whether selective depletion of CD38^+^ T cells can protect against diabetes onset *in vivo*, we treated eight weeks old diabetes-susceptible female NOD mice with weekly intraperitoneal injections of an anti-CD38 mAb (20 mg/kg) for eight weeks and monitored NBG for up to 40 weeks of age. Control mice received an equivalent volume of PBS for each corresponding injection. Five random mice from each group were sacrificed at week 20 for peri-islet infiltration analysis by immunohistochemistry (**Fig 3A**). In the control group, 19/24 (79%) mice developed hyperglycemia by ∼30 weeks, whereas only 5/28 (18%) mice in the anti-CD38 mAb treated group progressed to hyperglycemia by 40 weeks of age (**Fig 3B-C**). Furthermore, in a head-to-head comparison between anti-CD3 and anti-CD38 T cell depletion strategies, we treated 8 weeks old prediabetic, female NOD mice intraperitoneally injected with 50 μg of anti-CD3 F(ab)_2_ for 5 consecutive days or with a single dose of anti-CD38 mAb, analyzed the frequency of IGRP-specific autoreactive T cells on day 8 after the first injection (**Fig S3A-B**). Anti-CD3 F(ab)_2_ injection did not deplete IGRP-reactive CD8^+^ T cells compared to control (vehicle-injected) mice. Conversely, only one dose of anti-CD38 mAb injection led to a statistically significant decrease (*p*<0.001) in the percentage of IGRP-reactive CD8^+^ T cells compared to the anti-CD3 F(ab)_2_ treated group (**Fig S3C**). Anti-CD3 F(ab)_2_ did not confer durable protection against diabetes onset in NOD mice as indicated by only a transient delay in onset with no statistical difference compared to controls (**Fig 3D**). Kaplan-Meier analysis demonstrated significantly higher (*p*<0.0001) diabetes-free survival rate in the anti-CD38 mAb treated group compared to the control and CD3 F(ab)_2_ groups (**Fig 3E**). We then directly assessed peri-islet infiltration by harvesting pancreata at week 20 of age and performed H&E staining along with immunostaining for CD3 and CD38. H&E staining revealed a substantial peri-islet immune cell infiltration in the control mice compared to anti-CD38 mAb treated mice (**Fig 3F**). Immunohistochemistry data further confirmed substantially lower numbers of CD3^+^ and CD38^+^ cells within the islet microenvironment after anti-CD38 mAb treatment (**Fig 3G-H**). Collectively, these data indicated that selective ablation of CD38^+^ T cells conferred superior protection against autoimmune diabetes compared to broad CD3-directed depletion. These data implicate CD38^+^ autoreactive T cells as central drivers of islet autoimmunity and validate CD38 as a promising therapeutic target for T1D prevention in NOD mice.

**Fig 3.**
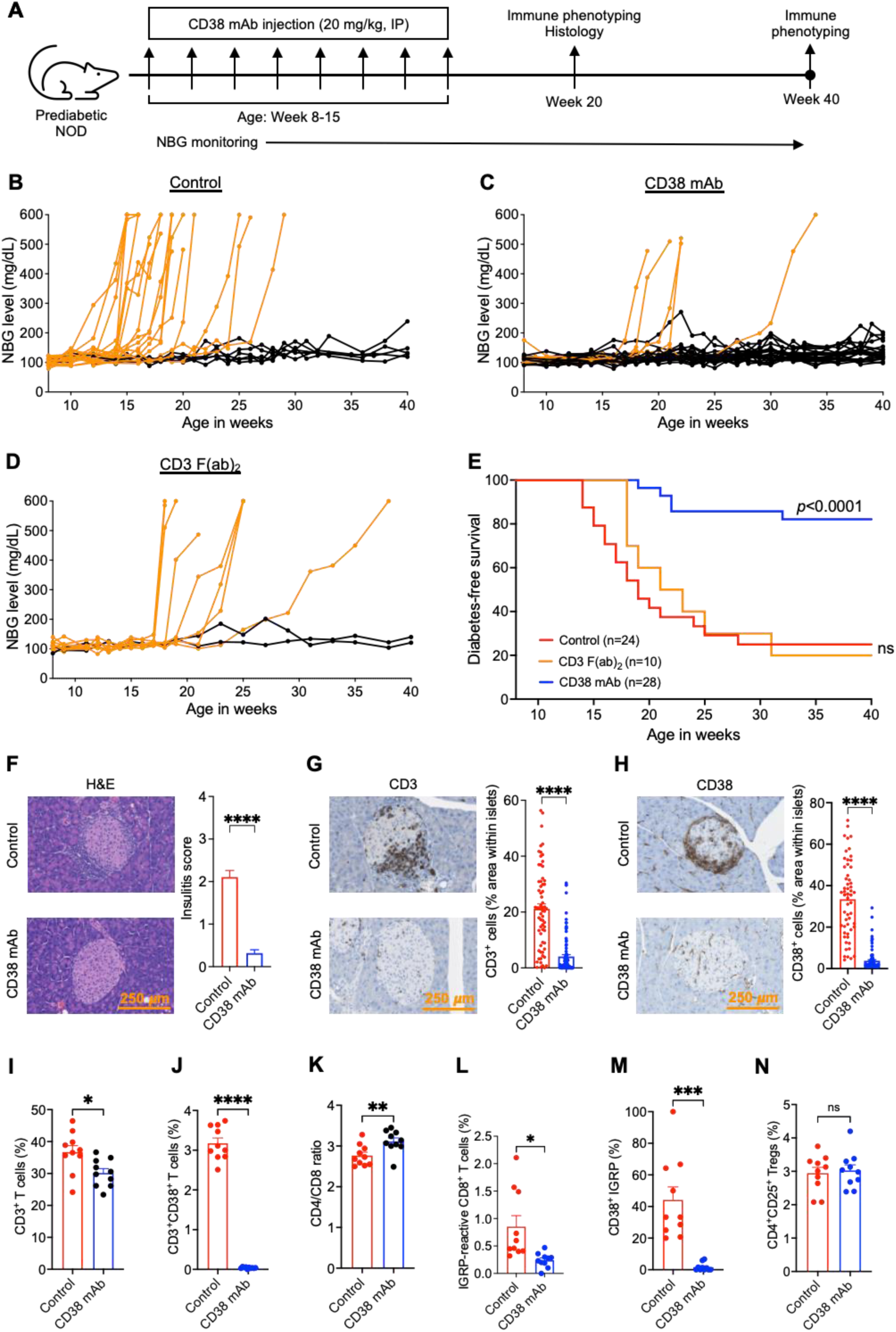
Anti-CD38 mAb prevents diabetes onset in NOD mice. **(A)** Eight weeks old female prediabetic NOD mice were treated weekly with intraperitoneal injections of anti-CD38 mAb at 20 mg/kg or with vehicle (control). In some experiments, NOD mice received 5 intravenous injections of anti-CD3 F(ab)_2_. Non-fasting blood glucose (NBG) was monitored weekly through week 40 of age. Some mice were sacrificed at week 20 for immune phenotyping and histological analyses. (**B**) NBG profiles of vehicle treated control mice. (**C**) NBG profiles of anti-CD38 mAb treated mice. (**D**) NBG profiles of anti-CD3 F(ab)_2_ treated mice. In **B-D**, each curve represents blood glucose profile of an individual mouse; orange curves denote mice that became hyperglycemic, whereas black curves denote mice that remained euglycemic. (**E**) Kaplan-Meier analysis of diabetes-free survival comparing control, anti-CD38 mAb treated, and anti-CD3 F(ab)_2_ treated mice. (**F**) Representative H&E-stained pancreas sections (left) and quantification of insulitis score (right) from control and anti-CD38 mAb treated NOD mice. Insulitis score calculated from 35-40 islets in each group. 0: no insulitis, 1: peri-insulitis, 2: infiltrative insulitis <50% of the islet, 3: infiltrative insulitis >50% of the islet. (**G-H**) CD3 and CD38 immunohistochemistry (IHC) of pancreas sections of control and anti-CD38 mAb treated NOD mice at 20 weeks of age. In **F**-**H**, Scale bar: 250 *µ*m; magnification: 40×. (**I-N**) Immunophenotyping of peripheral blood immune cell subsets in NOD mice at 20 weeks of age. Data represent mean±SEM and were pooled from at 2-4 independent experiments. ns: not significant; \**p*<0.05; \*\**p*<0.01; \*\*\**p*<0.001; \*\*\*\**p*<0.0001.

### Anti-CD38 mAb treatment leads to selective depletion of autoreactive T cells in NOD mice

To elucidate the mechanism by which anti-CD38 mAb abrogated autoimmunity, we performed multiparameter flow cytometric analysis of lymphocytes in anti-CD38 mAb treated NOD mice at weeks 20 of age and compared them with age- and sex-matched vehicle-treated controls. T cells were identified as CD3^+^, with CD4 and CD8 used to distinguish helper and cytotoxic T cell subsets, and CD38 used to distinguish CD38^+^ versus CD38^−^ cells. Anti-CD38 mAb caused only a modest reduction (*p*<0.05) in total CD3^+^ T cell frequency compared to control (**Fig 3I**). In contrast, depletion of CD38^+^ T cells was pronounced in anti-CD38 mAb treated NOD mice compared to control (*p*<0.0001) (**Fig 3J**). This shift produced an increased CD4/CD8 ratio in the anti-CD38 mAb treated group compared to control (*p*<0.01) (**Fig 3K**). Moreover, the frequency of IGRP-reactive CD8^+^ T cells was markedly reduced in anti-CD38 mAb treated group compared to control (*p*<0.05) (**Fig 3L**). Notably, the frequency of CD38^+^ IGRP-reactive CD8^+^ T cells was significantly lower in the anti-CD38 mAb treated group compared to control (*p*<0.001), indicating selective depletion of pathogenic autoreactive T cells and supporting a mechanism by which CD38-targeted therapy promotes long-term tolerance (**Fig 3M**). We also observed depletion of B220^+^ B cells with anti-CD38 mAb (*p*<0.001 *vs* control), with a more pronounced depletion within the CD38^+^ B cell subset (*p*<0.0001 *vs* control) (**Fig S4**). These findings suggest that anti-CD38 mAb may exert effects on both cellular and humoral compartment to promote tolerance. Importantly, anti-CD38 mAb treatment did not reduce the frequency of CD4^+^CD25^+^ Tregs, indicating preserved regulatory function (**Fig 3N**). The preferential preservation of Tregs with anti-CD38 mAb could enhance a tolerogenic milieu and contribute to durable immune tolerance in anti-CD38 mAb treated mice.

As anti-CD38 mAb conferred superior protection against autoimmune diabetes in NOD mice, we compared the lymphodepletion profiles of anti-CD38 mAb and anti-CD3 F(ab)_2_ in female NOD mice. Anti-CD38 mAb treatment was done on day 1 and day 8 (two doses) and anti-CD3 F(ab)_2_ treatment was done on day 1-5 (50 μg, 5 doses). We collected peripheral blood at day 14 and performed lymphodepletion analysis by flow cytometry (**Fig S5A-B**). Data showed that anti-CD38 mAb resulted in the milder global T-cell depletion than anti-CD3 F(ab)_2_ (*p*<0.05). Anti-CD38 mAb preferentially depleted CD8^+^ T cells which resulted in an increased CD4/CD8 T cell ratio. Further, IGRP-reactive CD8^+^ T cell depletion was more pronounced in anti-CD38 mAb treated group compared to CD3 F(ab)_2_ treated group (*p*<0.05 *vs* control) and this resulted in an increased CD4^+^CD25^+^ Treg/IGRP-reactive CD8^+^ T cell ratio, consistent with a more tolerogenic milieu (**Fig S5C**). Antigen-experienced T cells, including autoreactive clones, acquire a memory phenotype^15,16^. Thus, we investigated the effect of anti-CD38 mAb and anti-CD3 F(ab)_2_ on naïve and memory T cell compartment following lymphodepletion. Data showed that anti-CD38 mAb treatment induced minimal depletion of both CD4^+^ and CD8^+^ naïve (CD44^−^CD62L^+^) T cells, whereas anti-CD3 F(ab)_2_ caused pronounced naïve T cell depletion with a concomitant relative enrichment of T effector memory (TEM; CD44^+^CD62L^−^) and T central memory (TCM; CD44^+^CD62L^+^) T cells. In contrast, the anti-CD38 mAb treated group did not show a significant shift in the relative frequencies of both CD4^+^ and CD8^+^ TEM and TCM compared to control mice in peripheral blood (**Fig S6A-B**). Together with observed depletion of CD38^+^ T cells, these data demonstrated that anti-CD38 mAb selectively targets activated T cells, including islet antigen-specific T cells, while sparing naïve T cells.

To evaluate the durability of anti-CD38 mAb-mediated lymphodepletion effects, we performed longitudinal immune phenotyping in NOD mice at 40 weeks of age (**Fig S7A**). Because the control mice developed hyperglycemia early (median age of onset: 19 weeks), age-matched controls could not be maintained; accordingly, 20-week-old NOD mice were used as the controls for these studies. At 40 weeks of age, the overall T cell frequency and the CD4/CD8 ratio in the spleen did not differ significantly between anti-CD38 mAb treated mice and controls. Importantly, the frequency of IGRP-reactive CD8^+^ T cells was markedly lower in the anti-CD38 mAb treated group compared to control (*p*<0.01), indicating sustained suppression of autoreactive T cells. Notably, there was enrichment of CD4^+^CD25^+^ Tregs in the anti-CD38 mAb treated mice compared to controls (*p*<0.05) (**Fig S7B**), suggesting concomitant maintenance or expansion of the regulatory T cell compartment. Taken together, the concurrent enrichment of Treg subsets and the sustained reduction of autoreactive T cells support the establishment of peripheral tolerance and durable disease prevention in anti-CD38 mAb treated NOD mice.

### Anti-CD38 mAb treatment preserves islet integrity and function in NOD mice

T1D pathogenesis involves a bi-directional process in which autoreactive T cells infiltrate the pancreatic islets and islet-derived inflammatory signals amplify local immune destruction^17^. Senescent β-cells within the pancreatic islets of Langerhans contribute to this process by secreting senescence-associated secretory phenotype (SASP) factors, including pro-inflammatory cytokines, chemokines, and proteases, which can further impair islet function and promote inflammation^18,19^. These cells are characterized by established senescence markers including senescence-associated β-galactosidase (SA-β-Gal). To assess whether anti-CD38 mAb modulates islet biology beyond immune effects, we isolated islets from anti-CD38 mAb treated mice at 20 weeks of age and from age/sex-matched controls. We then performed absolute islet counting, and measured insulin content using N-(6-methoxy-8-quinolyl)-p-toluenesulfonamide) (TSQ) (a proxy for insulin granule abundance) (TSQ) and insulin immunohistochemistry. In addition, we assessed overall islet health by measuring oxygen consumption rate (OCR) and evaluating islet senescence (**Fig 4A**). Anti-CD38 mAb treated mice had higher number of islets compared to controls (*p*<0.001) (**Fig 4B**). TSQ fluorescence intensity was markedly increased, suggesting higher zinc content and preservation of zinc-rich insulin granules in the anti-CD38 mAb treated group (**Fig 4C-D**). To confirm that the TSQ staining reflected insulin content, we performed insulin immunohistochemistry on pancreas sections from control and anti-CD38 mAb treated mice. Control islets showed minimal insulin staining, whereas anti-CD38 mAb treated mice showed a robust insulin signal (*p*<0.001 *vs* control), indicating improved preservation of functional β-cells after anti-CD38 mAb treatment (**Fig 4E-F**). To assess islet metabolic function following anti-CD38 mAb treatment, we measured OCR in response to glucose stimulation. Islets from control mice exhibited a blunted OCR response to glucose, indicating impaired metabolic responsiveness. Moreover, islets from control mice showed minimal responses to oligomycin and carbonyl cyanide-4 (trifluoromethoxy) phenylhydrazone (FCCP), consistent with reduced ATP-linked and maximal respiratory capacity. In contrast, islets from anti-CD38 mAb treated mice showed robust glucose-stimulated increase in OCR, along with preserved responses to oligomycin and FCCP, indicating retained mitochondrial respiratory capacity (**Fig 4G**). Given that β-cells exhibit a senescence phenotype in T1D^20^, we next assessed whether anti-CD38 mAb treatment reduced islet senescence by staining dissociated islet cells with TSQ, CD38, 3H-Phenoxazin-3-one, 7-hydroxy-9H-9,9-dimethyl-, 10-oxide (DDAOG), and 5-Dodecanoylaminofluorescein Di-β-Galactopyranoside (C_12_FDG) (**Fig 4H**). Compared to control, islet cells from anti-CD38 mAb treated mice demonstrated decreased CD38 (*p*<0.01) and DDAOG (*p*<0.05) staining, indicating a low abundance of CD38-expressing and senescent β-cells (**Fig 4I-K, Fig S8).** The senescent cells showed co-staining of CD38 and DDAOG, and the treatment with anti-CD38 mAb decreased the percentage of CD38^+^DDAOG^+^ islets (**Fig 4I, 4L, Fig S9**). Furthermore, C_12_FDG staining of dissociated islets, a sensitive measure of SA-β-Gal, was significantly reduced in the islets from CD38 mAb-treated mice (*p*<0.05) (**Fig 4M-N**). Collectively, these data suggest that CD38-targeted therapy reduces the burden of senescent β-cells, promotes a less inflammatory environment, and restores β-cell function.

**Fig 4.**
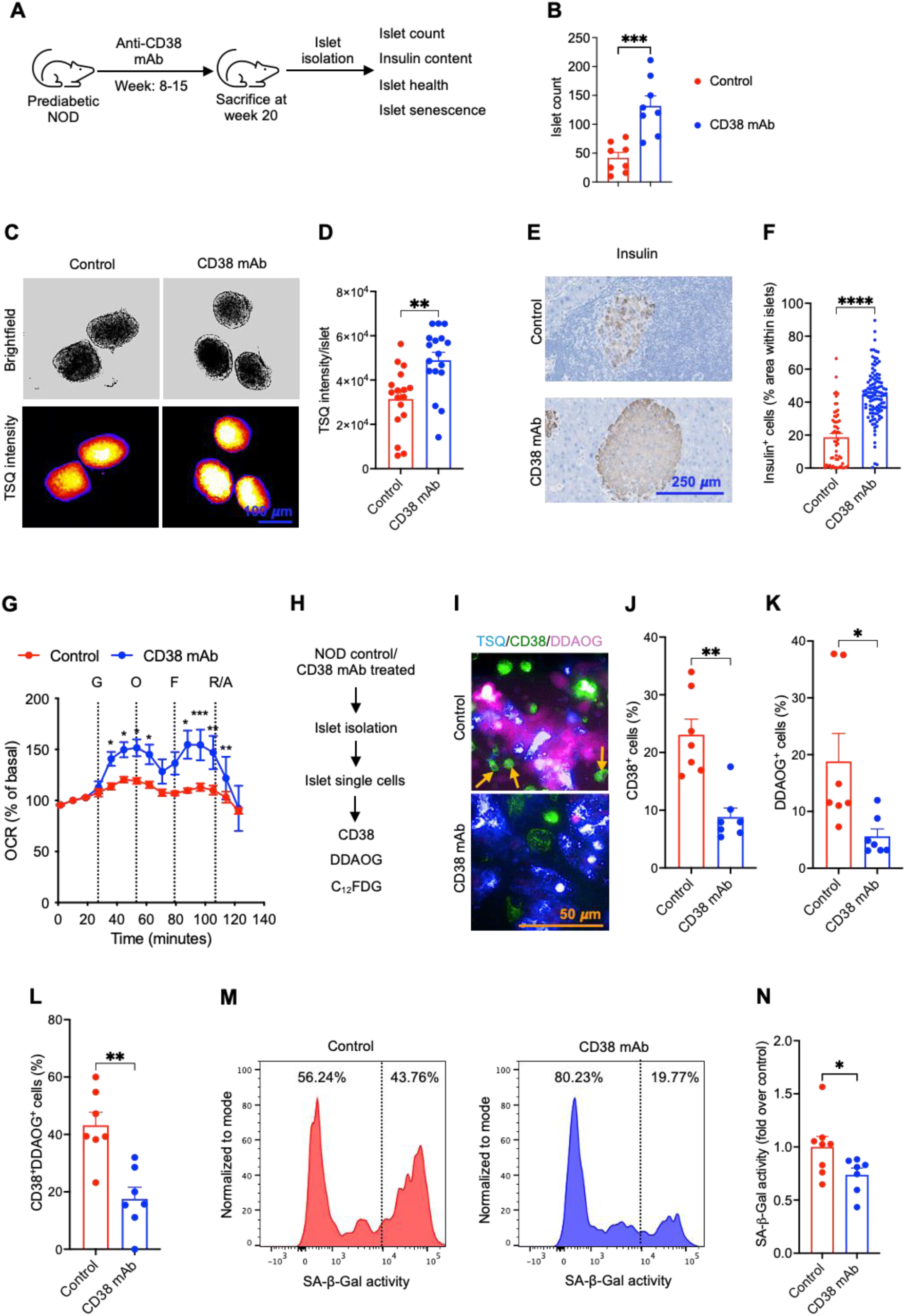
Anti-CD38 mAb treatment preserves islet integrity and function in NOD mice. **(A)** Eight weeks old female prediabetic NOD mice were treated weekly with intraperitoneal injections of anti-CD38 mAb at 20 mg/kg. Vehicle treated age- and sex-matched NOD mice were used as controls. Mice were sacrificed and analysis was performed at 20 weeks of age. (**B**) Number of islets isolated from control and anti-CD38 mAb treated mice. (**C**) Brightfield images (upper panel) and fluorescence images of intact islets in control and anti-CD38 mAb treated mice showing TSQ staining. (**D**) Quantification of the TSQ intensity in intact islets. (**E**) Representative image of insulin IHC staining. Magnification: 40×, Scale bar: 250 *μ*m. (**F**) Quantification of insulin IHC. (**G**) Measurement of OCR upon sequential injections of 16.7 mM glucose (G), oligomycin (O), FCCP (F), and rotenone/antimycin A (R/A). (**H**) Design of experiment to evaluate islet integrity and function in dissociated cells. (**I**) Representative fluorescence images of islet single cells showing TSQ, DDAOG, and CD38 staining. Arrows indicate CD38^+^DDAOG**^+^** cells. Magnification: 40×, Scale bar: 100 *μ*m. Quantification of fluorescence intensity of (**J**) CD38, (**K**) DDAOG, and (**L**) CD38^+^DDAOG^+^ staining. (**M**) Representative histograms showing SA-β-Gal activity in control and anti-CD38 mAb treated mice. (**N**) Quantification of the SA-β-Gal activity. Data represent mean±SEM and were pooled from three independent experiments. ns: not significant; \**p*<0.05; \*\**p*<0.01; \*\*\**p*<0.001; \*\*\*\**p*<0.0001.

### Circulating CD38^+^ autoreactive T cells are present at T1D diagnosis but decrease in established T1D patients

To assess the clinical relevance of the murine CD38-driven T1D pathogenesis observed in diabetes-susceptible NOD mice, we analyzed peripheral blood mononuclear cells (PBMCs) from three Human Leukocyte Antigen-A2 positive (HLA-A2^+^) cohorts: healthy donors, recently-diagnosed T1D, and individuals with established T1D (onset >1 year) (**Fig 5A**). We generated HLA-A2-restricted p-HLA multimers (spheromers) presenting GAD65 (p110-119; FLQDVMNILL) to detect autoreactive T cells among human patient samples (**Fig 5B, S10**). PBMCs were stained with p-HLA-A2 spheromers conjugated with AF647 and PE to analyze the frequency of autoreactive CD8^+^ T cells by flow cytometry. We co-stained with CD38 antibody to identify and track expression of CD38 among autoreactive T cells. Spheromer-bound CD8^+^ T cells were detected across all cohorts, indicating the presence of autoreactive T cells irrespective of disease stage. These findings align with prior work that reported detectable autoreactive T cells in both healthy donors and individuals with T1D without a statistically significant difference between groups, confirming that autoreactive T cells are present in healthy individuals^21^.

**Fig 5.**
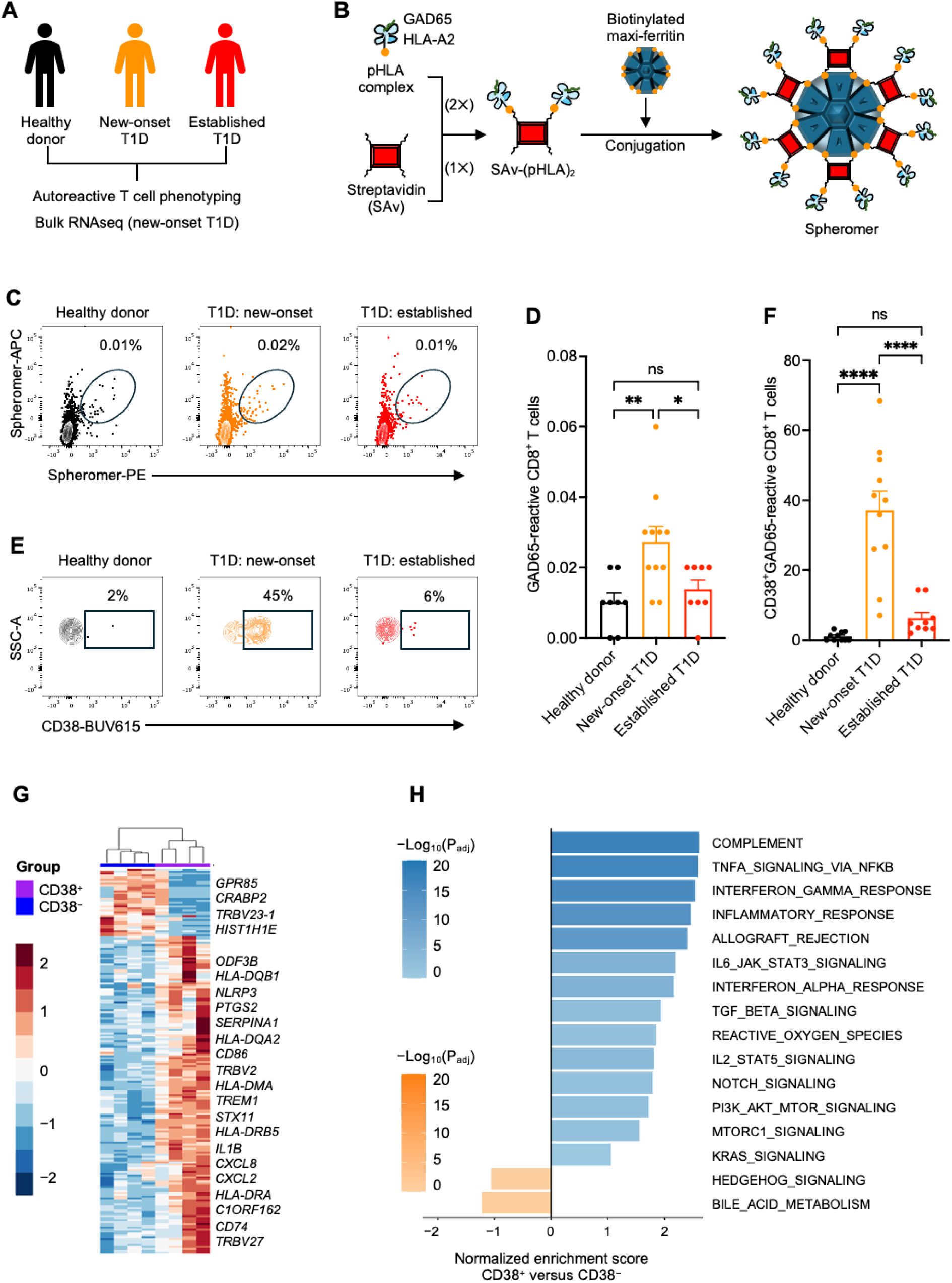
CD38^+^ autoreactive T cells found in recently diagnosed T1D patients are phenotypically distinct from CD38^−^ autoreactive T cells. **(A)** Peripheral blood mononuclear cells (PBMCs) were collected from healthy donors, patients with new-onset T1D, and patients with established T1D, and CD38 expression was assessed among the autoreactive T cells by flow cytometry. (**B**) Assembly workflow of HLA-A2 restricted GAD65-specific spheromer reagents. Biotinylated peptide-HLA-A02:01 (pHLA) monomers containing GAD65 (p110-119) was first complexed with streptavidin (SAv) to generate intermediate SAv-(pHLA)₂ complexes. These intermediates were subsequently conjugated to a biotinylated 24-mer maxi-ferritin nanoparticle scaffold to assemble multivalent spheromer complexes. Purified spheromer reagents were used for downstream staining and phenotypic analysis of GAD65-reactive CD8^+^ T cells. (**C**) Representative flow cytometry scatter plots showing frequency of GAD65-reactive CD8^+^ T cells in healthy donors, new-onset T1D patients, and established T1D patients. (**D**) Quantification of GAD65-reactive CD8^+^ T cells across groups. (**E**) Representative flow cytometry contour plots showing surface expression of CD38 among GAD65-reactive CD8^+^ T cells. (**F**) Quantification of CD38-expressing GAD65-reactive CD8^+^ T cells across groups. (**G-H**) Bulk RNA sequencing of CD38^+^ and CD38^−^ autoreactive T cells. For bulk RNA sequencing of CD38^+^ and CD38^−^autoreactive T cells, GAD65- and IGRP-reactive CD8^+^ T cells were pooled. (**G**) Heatmap and hierarchical clustering of top 200 most differentially expressed genes by adjusted *p*-values in CD38^+^ versus CD38^−^ autoreactive T cells. (**H**) Gene-set enrichment analysis of CD38^+^ versus CD38^−^ autoreactive T cells. The 16 most immune-related and differentially expressed pathways by normalized enrichment score are shown. ns: not significant; \**p*<0.05; \*\**p*<0.01; \*\*\**p*<0.001; \*\*\*\**p*<0.0001.

However, in a stratified analysis of T1D patient samples, we observed that the percentage of GAD65-reactive CD8^+^ T cells was significantly higher in new-onset patients compared to healthy controls (*p*<0.01) or established T1D patients (*p*<0.05) (**Fig 5C-D**). This pattern shows that autoreactive T cell expansion is most readily detected during disease onset and wanes with disease progression. Notably, patients recently diagnosed with T1D were characterized by higher CD38 expression among GAD65-specific autoreactive CD8^+^ T cells when compared to healthy controls and individuals with established T1D across three independent cohorts (*p*<0.0001 *vs* healthy donor or established T1D) (**Fig 5E-F**). Additionally, IGRP (p227-236; VLNIDLLWSV)-reactive CD8^+^ T cells from recently diagnosed T1D patients showed a similar phenotype, with a significantly higher fraction of CD38^+^ autoreactive T cells compared to healthy donors or established T1D (**Fig S11**). Given the similar phenotypes of GAD65- and IGRP-reactive CD8^+^ T cells, our data support hypothesis that CD38 represents a specific marker of pathogenic CD8^+^ T cells targeting β-cell antigens among T1D patients during the time of clinical diagnosis.

### CD38 expression defines a unique transcriptional signature in autoreactive T cells

To further characterize the human autoreactive T cell subsets, we sorted CD38^-^ and CD38^+^ autoreactive T cells and performed bulk RNAseq transcriptomics analysis. Spheromers generated separately using IGRP and GAD65 peptides were pooled and used to stain PBMCs to improve sort yield from recently diagnosed T1D patients (**Fig S12A**). Principal component (PC) analysis revealed that CD38^+^ autoreactive T cells formed a transcriptional cluster with high intra-group consistency, that was distinct from CD38^−^ autoreactive T cells (**Fig S12B**). Differential expression analysis identified 391 upregulated and 158 downregulated genes in CD38^+^ autoreactive T cells *versus* CD38^−^ autoreactive T cells (**Fig S12C**). Hierarchical clustering and heatmap analysis of CD38^+^ *versus* CD38^−^ autoreactive T cells revealed a broad transcriptional reprogramming towards an activated, proinflammatory effector phenotype. Multiple HLA genes including *HLA-DMA*, *HLA-DQA2*, *HLA-DQB1*, *HLA-DR*, *HLA-DRA*, and *HLA-DRB5* were markedly upregulated in CD38^+^ autoreactive T cells, suggesting an activated state of CD38^+^ autoreactive T cells. Concomitant induction of inflammatory mediators such as CXCL2, *CXCL8*, *IL1B*, *NLRP3, PTGS2*, and *TREM1* indicated activation of innate-like inflammasome and chemokine pathways in CD38^+^ autoreactive T cells. Upregulation of activation markers such as *CD86*, *PSMB10,* and *STX11* suggested amplified cytolytic potential and metabolic adaptation to sustained immune activation. Notably, selective enrichment of TCR-B genes including *TRBV2*, *TRBV7-3*, *TRBV27*, *TRBJ2-6* suggested oligoclonal expansion of CD38^+^ autoreactive CD8^+^ T cell clones possibly driven by β-cell derived epitopes (**Fig 5G, S12D**). These transcriptional features of CD38^+^ autoreactive T cells delineate a pathogenic CD8^+^ T cell population exhibiting proinflammatory responses among patients recently diagnosed with T1D. Furthermore, gene-set enrichment analysis of CD38^+^ *versus* CD38^−^autoreactive T cells revealed a significant upregulation of inflammatory, interferon, TNF/NF-kB, IL6-JAK-STAT3, Notch, complement, and allograft rejection-related signatures in CD38^+^ autoreactive T cells. This was accompanied by enrichment of IL2-STAT5, PI3K-AKT-mTOR, and MTORC1 pathways, consistent with enhanced survival, proliferation, and metabolic reprogramming of CD38^+^ compared to CD38^-^ autoreactive T cells (**Fig 5H**). Together, these data suggest that CD38^+^ autoreactive T cells are transcriptionally distinct and help drive β-cell autoimmunity compared to CD38^−^ autoreactive T cells.

### Human anti-CD38 mAb prevents insulitis in a humanized model of T1D

To verify whether CD38^+^ autoreactive T cell drive human autoimmune diabetes pathogenesis, we conducted a series of experiments wherein PBMCs from recently diagnosed HLA-A2^+^ T1D patients were adoptively transferred into NSG-HLA-A2/HHD mice. Following PBMC injection, recipients were randomized to receive weekly doses of anti-human CD38 mAb or PBS as controls (**Fig 6A**). Blood glucose levels of mice in both groups remained in the normal range throughout the study period (**Fig S13**). However, when pancreata were harvested and evaluated by H&E staining at day 35 post-transfer, mice that received PBMCs alone exhibited extensive peri-islet infiltration (insulitis), whereas anti-CD38 mAb treated mice showed markedly reduced peri-islet infiltration, indicating protection from infiltrating CD38^+^ pathogenic T cells (**Fig 6B-C**). We harvested spleens and performed flow cytometric analysis to evaluate the lymphodepletion effect of anti-CD38 mAb. The frequency of total CD3^+^ T cells in anti-CD38 mAb treated group was marginally decreased compared to PBMC-only group (*p*<0.05) (**Fig 6D**). Approximately 30% of CD3^+^ T cells in the PBMC-only group expressed CD38, whereas treatment with anti-CD38 mAb virtually eliminated these cells (*p*<0.01) (**Fig 6E**). Notably, the proportion of CD38^+^ GAD65-reactive CD8^+^ T cells was significantly lower in the anti-CD38 mAb treated group compared to PBMC-only group (*p*<0.0001) (**Fig 6F**). Taken together, these data support the premise that CD38^+^ autoreactive T cells contribute to disease progression in mice and humans with T1D. Selective depletion of CD38^+^ autoreactive T cells mitigates islet inflammatory infiltration in this humanized T1D model.

**Fig. 6.**
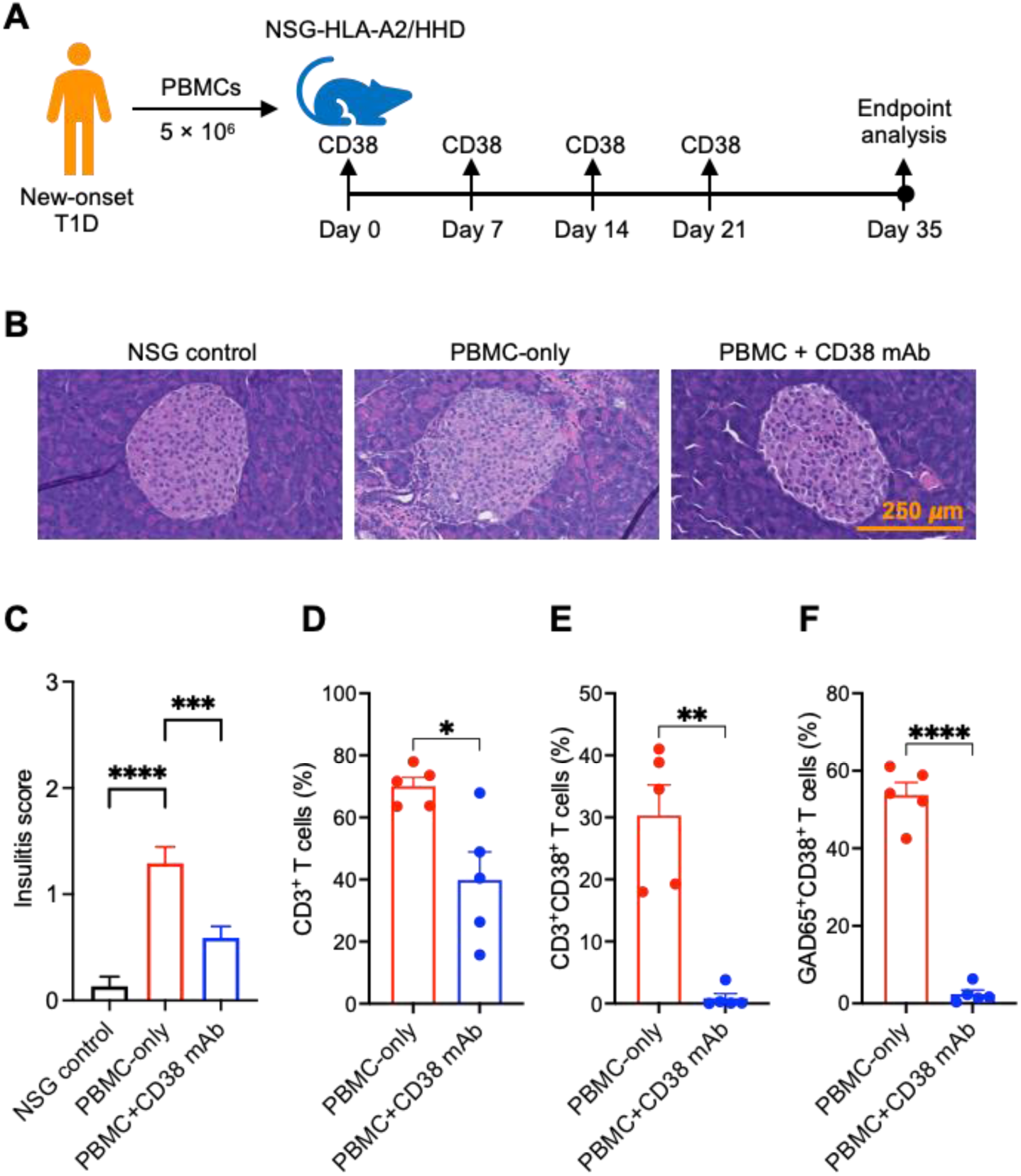
Anti-CD38 mAb prevents insulitis in a humanized mouse model of T1D. (**A**) Design of the *in vivo* proof-of-concept experiment to assess the efficacy of anti-human CD38 mAb in a humanized mouse model of T1D. PBMCs from new-onset T1D patients were transferred into NSG-HLA-A2/HHD mice. Mice received weekly injections of anti-CD38 mAb for four weeks; PBMC-alone mice served as controls. Mice were sacrificed at day 35 post-transfer for histological and lymphodepletion analyses. (**B**) Representative H&E images showing peri-islet lymphocytic infiltration in NSG-HLA-A2/HHD recipients. Magnification: 40’, Scale bar: 250 *m*m. (**C**) Insulitis score calculated from 35 islets in each group. 0: no insulitis, 1: peri-insulitis, 2: infiltrative insulitis <50% of the islet, 3: infiltrative insulitis >50% of the islet. (**D**) Percentage of CD3^+^ T cells in spleen. (**E**) Percentage of CD3^+^CD38^+^ T cells in spleen. (**F**) Percentage of CD38^+^ GAD65-reactive CD8^+^ T cells in spleen. Data were pooled from two independent experiments. ns: not significant, \**p*<0.05, \*\**p*<0.01, \*\*\**p*<0.001, \*\*\*\**p*<0.0001.

## DISCUSSION

The currently approved anti-CD3 mAb for T1D, teplizumab, transiently delays T1D progression while indiscriminately depleting both pathogenic effector T cells and Tregs. This non-specific depleting activity might inhibit its effectiveness in establishing durable tolerance^4,10,11^. Furthermore, patients undergoing long-term treatment with teplizumab may be at risk for serious infections due to prolonged lymphopenia^22^. In this study, we identified that CD38 expression was higher on pathogenic autoreactive T cells in diabetes-susceptible mice and humans with T1D. Our findings further demonstrate that anti-CD38 mAb effectively mitigates T1D progression in murine and humanized models, by selectively depleting CD38^+^ diabetogenic T cells.

Autoreactive T cells are defined not only by antigen specificity but also by a distinct, disease-imprinted cellular state^23,24^. In this study, IGRP-reactive CD8^+^ T cells exhibit a unique clonal expansion and a transcriptional program exhibiting activation, proliferation, and inflammatory pathways. In contrast, non-autoreactive T cells show quiescent or less pathogenic states as indicated by downregulated activation markers at transcriptomics level. Conceptually, this autoreactive state shares features with chronically stimulated malignant lymphocytes or plasma cells, wherein sustained signaling and metabolic reprogramming define pathogenic clones. Markers like CD38 reflect this activated, metabolically reprogrammed phenotype^25^. A striking finding of the current study is that among human GAD65 and IGRP autoreactive T cells that overexpress CD38 show a greatly enriched inflammatory and proliferative phenotype compared to CD38^−^ autoreactive T cells. Further evaluation is needed to better understand the differences between these subgroups of autoreactive T cells, as well as how CD38^−^ autoreactive T cells might indirectly or actively contribute to immune tolerance after *in vivo* anti-CD38 mAb depletion.

The clinical efficacy of CD38 targeting antibodies has been primarily studied for the treatment of plasma cell dyscrasias, where the excessive production of non-functional autoantibodies can lead to direct organ toxicity and paraneoplastic autoimmune disorders^26,27^. Monoclonal antibodies like daratumumab and isatuximab are FDA approved and have become part of the standard backbone treatment for multiple myeloma due to their overall survival benefit^26,27^. Given their favorable safety profile and efficacy, the use of these antibodies extends beyond the treatment of hematological malignancies. Daratumumab is now used for the treatment of refractory immune-mediated thrombocytopenia^28^, autoimmune hemolytic anemia^29^, pure red cell aplasia due to ABO mismatch^30^, and desensitization of patients with anti-donor antibodies in the setting of transplantation^31^. While T1D is commonly considered to be driven by autoreactive T cells, the disease has accompanied robust antibody recognition of islet-specific antigens, and the humoral responses undoubtedly contribute. It is likely that CD38 depletion also reduces pathogenic plasmablasts and plasma cells, suggesting that anti-CD38 mAb therapy could be synergistic. More broadly, defining a deep phenotype of CD38-dependent disease-causing autoreactive T or B cells across autoimmune diseases beyond T1D can enhance mechanistic understanding and guide development of targeted therapies that selectively eliminate or reprogram pathogenic clones while maintaining the protective immunity.

While anti-CD38 mAb has been broadly used across various disease conditions, individuals with T1D include pediatric or young adults and long-term safety remains an important consideration including potential off-target effects, and effects on humoral immunity and B cell function. Nevertheless, there is a strong rationale for evaluating anti-CD38 mAb therapy in clinical trials in adults with recent-onset T1D to evaluate safety and tolerability, C-peptide preservation, delay in insulin independence, and immune correlates. Unlike anti-CD3 treatment, anti-CD38 mAb therapy could be easily integrated into combination strategies with antigen-specific therapies or approaches to rebalance tolerance pathways including low-dose IL-2, or Treg cell therapies. Anti-CD38 mAb therapy may also provide a robust target for B and T cell depletion strategies in T1D utilizing chimeric antigen receptors or bispecifics which have shown increased efficacy in other contexts^32,33^. Our data suggest that the feasibility of anti-CD38 mAb therapy could extend beyond T1D and in other T cell driven immune dysregulation, as CD38 overexpression was observed in activated T cells in graft versus host disease patients (**Fig S14**).

The major limitations of this study should be acknowledged. Although we identified a marked upregulation of CD38 in autoreactive T cells from patients recently diagnosed with T1D, the mechanism underlying the apparent loss of this population in longstanding disease remains to be investigated. It is unclear whether autoreactive T cells exhibit plasticity between CD38^+^ pathogenic and CD38^-^ non-pathogenic states depending on the stage of disease. In addition, our *in vivo* analysis was limited to a single dose (20 mg/kg) of anti-CD38 mAb in the murine T1D model, which was effective in preventing disease; however, rigorous dose-escalation studies will be required to define the optimal efficacious dose and therapeutic window. Future work should also assess combination strategies using lower doses of anti-CD38 mAb together with other immunomodulatory agents to maximize the therapeutic efficacy and minimize adverse and off-target effects. Further, the long-term impact of anti-CD38 mAb treatment on other immune compartments, including B cells, dendritic cells, and natural killer cells, remains to be determined and will be essential to fully understand the mechanisms that underlie durable immune tolerance in T1D. Finally, whether CD38^+^ autoreactive T cells contribute to senescence-associated remodeling of the islet microenvironment or participate in broader inflammatory circuits that sustain disease progression warrants future investigation.

In conclusion, CD38 represents a compelling, mechanistically plausible target for selectively depleting autoreactive T cells that drive T1D pathogenesis. The dual utility of CD38 as a functional modulator of autoreactive T cells and as a disease-relevant biomarker offers a two-pronged strategy: (1) enabling earlier, more precise disease detection and monitoring of T1D and (2) preventing the autoimmune attack on β-cells by allowing a selective depletion of CD38^+^ pathogenic autoreactive T cells. With rational patient selection, optimized dosing, and rigorous biomarker validation, CD38-targeted therapies could complement existing preventive and disease-modifying approaches. The strategy of selectively targeting CD38 could redefine the therapeutic landscape of T1D, shifting the paradigm to proactive immune modulation at a stage when the disease is most amenable to preservation of endogenous β-cell mass.

## MATERIALS AND METHODS

### Mice

All experimental mice were purchased from the Jackson Laboratory (Sacramento, CA) and housed in the Stanford University Animal Facility. Animal experiments were followed according to the protocol approved by Institutional Animal Care and Use Committee of Stanford University (IACUC: 30219). Prediabetic female NOD/ShiLtJ mice (stock no. 001976) were used for diabetes onset study. For adoptive T cell transfer experiments, NOD-Rag1^null^IL2rg^null^ (Rag1-KO/NRG; stock no. 007799) mice were used. For humanized T1D model, NOD.Cg-Prkdc^scid^ Il2rg^tm1Wjl^ Tg (HLA-A/H2-D/B2M)1Dvs/SzJ (NSG-HLA-A2/HHD; stock no. 014570) mice were used.

### Human samples

Individuals with T1D provided informed consent in accordance with the Declaration of Helsinki and were enrolled in a research protocol approved by the Stanford University Institutional Review Board (IRB) protocol #35453. Blood sample was collected, and peripheral blood mononuclear cells (PBMCs) were isolated by density gradient centrifugation using Ficoll-Paque (Cytiva, Marlborough, MA). For healthy controls, blood samples were obtained from Stanford Blood Center (Palo Alto, CA) and PBMCs were isolated.

### Detection of autoreactive T cells in NOD mice

Islet-specific glucose-6-phosphatase subunit-related protein (IGRP) specific CD8^+^ T cells in NOD mice were identified using tetramers provided from the NIH Tetramer Core Facility. Cells were Fc receptor blocked and stained with APC- and PE-conjugated IGRP tetramers (Sequence 206-214: VYLKTNVFL). Tetramers were used at a dilution of 1:200 for 30 min at 4°C. Cells were washed after tetramer staining and additionally stained with antibodies for flow cytometry.

### Generation of pHLA-spheromer reagents

HLA-A*02:01-restricted peptide epitopes derived from the T1D autoantigens GAD65 (p110-119; FLQDVMNILL) and IGRP (p227-236; VLNIDLLWSV) were selected and evaluated for predicted HLA-A2 binding affinity using the Immune Epitope Database (IEDB) analysis platform. We used cancer/testis antigen 83 (p90-98; KLVELEHTL) as a non-specific antigen for spheromer preparation. The synthetic peptides (Peptide 2.0 Inc.) were used for *in vitro* refolding with recombinant HLA-A*02:01 heavy chain and β2-microglobulin (β2m), which were expressed in E. coli as inclusion bodies following established protocols from the NIH Tetramer Core Facility. Briefly, denatured HLA-A*02:01 heavy chain and β2m were refolded in the presence of individual peptide at 4°C for 72-96 hours. Refolded peptide-HLA (pHLA) complexes were purified by size-exclusion chromatography (SEC), and successful complex formation was confirmed based on SEC elution profiles and SDS-PAGE analysis (**Fig S10**). Purified pHLA monomers were biotinylated using BirA biotin ligase and subsequently assembled into multivalent spheromer reagents using a two-step biotin-streptavidin conjugation strategy with a biotinylated 24-mer maxi-ferritin nanoparticle scaffold, as previously described^34^. Briefly, biotinylated pHLA monomers were first complexed with streptavidin to generate intermediate SAv-(pHLA)₂ complexes, followed by conjugation to the biotinylated nanoparticle scaffold. Assembled spheromer complexes were purified by SEC and used for downstream staining and phenotypic analysis of antigen-specific CD8 T cells.

### Detection of human autoreactive T cells

The pHLA monomers were assembled into spheromer complexes conjugated with either PE or AF647 fluorophores using streptavidin-based assembly. For flow cytometric analysis, PBMCs were stained with PE- and APC-conjugated GAD65 or IGRP spheromers. For sorting of autoreactive T cells, GAD65 and IGRP spheromers were pooled to generate a staining cocktail. Cryopreserved PBMC samples were thawed and incubated with Fc receptor blocking reagent to minimize non-specific binding with antibodies and spheromers. Cells were stained with pooled PE- and AF647-labeled spheromer reagents in staining buffer for 1 hour at 4°C in the dark. Following spheromer staining, cells were washed and stained with antibodies against surface markers, including CD3, CD8, and CD38, along with a viability dye to exclude dead cells. Stained samples were sorted using a fluorescence-activated cell sorting (FACS) instrument (FACSAria Fusion). Antigen-specific CD8^+^ T cells were identified by dual-positive spheromer staining (PE^+^AF647^+^) within live CD3^+^CD8^+^ lymphocytes. Spheromer^+^CD8 T cells were further separated into CD38^+^ and CD38^-^ subsets based on surface CD38 expression. Sorted populations were collected directly into the appropriate collection buffer and subsequently used for bulk RNA-sequencing library preparation and downstream transcriptomic analysis.

### Flow cytometry

Live cells were identified by Live/Dead Aqua (BioLegend; Cat. 423102) and stained with antibodies at a final dilution of 1:200. The following antibodies were used for flow cytometry analysis in mice: Brilliant Ultraviolet 737 (BUV737)-conjugated CD3 (BD Biosciences; Cat. 612803), BUV395-conjugated CD4 (Invitrogen; Cat. 363-0042-82), BUV805-conjugated CD8 (BD Biosciences; Cat. 612898), Allophycocyanin Cyanine 7 (APC/Cy7)-conjugated B220 (BioLegend; Cat. 103224), BV711-conjugated CD25 (BioLegend; Cat. 102049), Alexa Flour 700 (AF700)-conjugated CD44 (BioLegend; Cat. 103026), Brilliant Violet 605 (BV605)-conjugated CD62L (BioLegend; Cat. 104438), Fluorescein isothiocyanate (FITC)-conjugated CD38 (BioLegend; Cat. 102705), PB-conjugated Foxp3 (BioLegend, Cat. 126410). The following antibodies were used for staining human samples: FITC-conjugated CD3 (Biolegend; Cat. 317306), BV605-conjugated CD19 (Biolegend; Cat. 302244), BV711-conjugated CD4 (Biolegend; Cat. 317440), APC/CY7-conjugated CD8 (Biolegend; Cat. 344714), and BV421-conjugated CD38 (Biolegend; Cat. 303526). Data were collected using BD FACSymphony flow cytometer and analyzed by FlowJo Software Version 10.10.1 (Tree Star, Ashland, OR).

### Bulk RNA sequencing

Total RNA was isolated using NucleoSpin RNA Kit (Ref: 740955.50) and quantified using a Qubit Fluorometer (Invitrogen) and TapeStation 4200 (Agilent Technologies) for quality assessment. RNA libraries were prepared using the Takara SMART-Seq mRNA Kit (Cat. 634773) following the manufacturer’s instructions. This workflow employs SMART (Switching Mechanism at the 5′ end of RNA Template) technology to generate full-length cDNA from polyadenylated RNA transcripts. First-strand cDNA synthesis and template switching were followed by LD PCR amplification to enrich for full-length cDNA molecules. The amplified cDNA was purified using Agencourt AMPure XP beads (Beckman Coulter), and library quality and quantity were confirmed using TapeStation 4200 and Qubit. Libraries were constructed using the Nextera XT DNA Library Preparation Kit (Illumina, Cat. FC-131-1096) and indexed *via* tagmentation and PCR amplification to generate Illumina-compatible libraries. Indexed libraries were pooled, normalized to 4 nM, and sequenced on an Illumina NovaSeq X Plus platform using paired-end (2×100 bp) reads.

### Adoptive T cell transfer

Diabetes-susceptible NOD mice (12 weeks old) were used as T cell donors. CD3^+^ T cells (±CD38^+^ T cells) were sorted to purity using BD FACSAria II at Stanford Shared FACS facility. Then, 3 × 10^6^ CD3^+^ T cells were intravenously injected into each NRG recipient. The recipient NRG mice were monitored for diabetes induction by weekly checking their non-fasting blood glucose levels using Contour Next EZ blood glucose monitoring system (Ascensia Diabetes Care, Parsippany, NJ). The mice were considered hyperglycemic when two consecutive blood glucose measurements exceeded 400 mg/dL. Peripheral blood sample was collected at day 20 to analyze lymphocyte expansion *in vivo*.

### Diabetes onset study

Eight weeks old female prediabetic NOD mice were treated with eight weekly doses of 20 mg/kg of anti-CD38 antibody *via* intraperitoneal injection (BioXcell; BE0317). Dose of anti-CD3 F(ab)_2_ (BioXcell; BE0001-1FAB) was selected based on previously published data^35^ and was injected intraperitoneally. Non-fasting blood glucose was monitored weekly until week 40. The mice were considered hyperglycemic when two consecutive blood glucose measurements exceeded 400 mg/dL.

### Histological Analysis

Mice were euthanized, and pancreata were isolated and fixed in formalin before being embedded in paraffin blocks. Pancreas sections were stained with hematoxylin and eosin using the method described previously^36^. Immunohistochemical staining was performed using methods reported previously^36–38^. Insulin (Cell Signaling, clone C27C9, dilution 1:200), anti-CD3 (Cell Signaling, clone D7A6E, dilution 1:200) and anti-CD38 (Cell Signaling, clone E9F5A, dilution 1:150) were used for immunohistochemistry analysis.

### Fluorescent staining of pancreatic islets

Pancreatic islets were isolated using a method described previously^39^. Isolated islets were cultured overnight in DMEM/low glucose medium (Cytiva) containing 10% fetal bovine serum (Gibco) and 1% penicillin/streptomycin (Gibco). Islets were incubated with 10 μM of (N-(6-methoxy-8-quinolyl)-p-toluenesulfonamide) (TSQ) for 30 min at 37° C. For dissociated islet staining, islets were first dissociated with TrypLE Express (Thermo Fisher Scientific), plated into 384-well plates precoated with conditioned media from 804G rat bladder carcinoma cells and allowed to attach for 24 hour before incubating with 10 μM TSQ, FITC-conjugated CD38 (1:100) and 10 μM 9H-(1,3-Dichloro-9,9-Dimethylacridin-2-One-7-yl) β-D-Galactopyranoside (DDAOG; Invitrogen) for 45 min at 37° C. Staining was imaged on the Operetta CLS (Revvity) in the DAPI (TSQ), FITC (CD38) and Cy5 (DDAOG) channel. Fluorescence intensity was quantified on the Harmony Software Version 5.2 (Revvity).

### Assessment of islet senescence

Isolated islets were purified by handpicking and dissociated into single cells and treated with 100 nM Bafilomycin A1 for 1 hour. Islet single cells were stained with 33 μM of 5-dodecanoylaminofluorescein di-β-D-galactopyranoside (C_12_FDG; Invitrogen) for 1 hour at 37° C. Cells were then stained with viability dye (Live/Dead Aqua) and BV711-conjugated CD45 antibody to exclude hematopoietic lineage cells. Senescence-associated β galactosidase (SA-β-Gal) activity was assessed by flow cytometry using the 488 nm laser line.

### Seahorse extracellular flux assay

Respirometry measurements were performed using a Seahorse XFe24 Analyzer (Agilent Technologies) according to the manufacturer’s instructions. Briefly, ∼50 islets were seeded in Islet Capture Microplates and incubated in Seahorse XF Media (3 mmol/L glucose and 1% FBS) for 1 hour in a CO_2_-free 37° C incubator, and oxygen consumption rate (OCR) was measured upon sequential injections of 16.7 mM glucose, 5 μM oligomycin, 3 µM carbonyl cyanide-4 (trifluoromethoxy) phenylhydrazone (FCCP), and 5 µM rotenone/antimycin A (Agilent Technologies).

### Humanized T1D model

PBMCs from new-onset T1D patients were intravenously injected into 8 weeks old female NSG-HLA-A2/HHD mice. Each mouse received 5 × 10^6^ PBMCs. Following PBMC injection, some mice received weekly intraperitoneal injections of human anti-CD38 mAb (BioXcell; SIM0034) at a dose of 20 mg/kg for 4 weeks. Mice were euthanized at day 35 following PBMC injection and histological analysis of pancreas sections was performed using hematoxylin and eosin staining.

### Statistical Analysis

Statistical analysis was performed by using GraphPad Prism 11 (La Jolla, CA). Statistical values were calculated using unpaired *t*-test or one-way ANOVA. Differences with *p* values less than 0.05 were considered statistically significant.

## Acknowledgments

We thank the staff of the Stanford University Animal Care Facility for taking exceptional care of the mice during the experiments. We also thank the NIH Tetramer Core Facility for generously providing tetramers. We are deeply grateful to the patients who generously donated blood samples for this study; their participation was essential to the conduct and success of this work.

## Author contributions

Conceptualization: SP, EHM

Methodology: SP, SL, SR, CSB, NN, KPJ, EHM. RA developed spheromers for autoreactive T cell identification in human T1D samples.

Investigation: SP, RA, SL, CSB, SR, BPI, PC, BG, AP, YBK, NH, KPJ, PJU, NN, MMD, JPA, EHM

Visualization: SP, RA, SL, SR, CSB, NN, MMD, JPA, EHM

Funding acquisition: EHM, AST

Project administration: NN, KPJ

Supervision: EHM

Writing – original draft: SP

Feedback on original draft: RA, SL, CSB, SR, BPI, PC, BG, AP, NH, KPJ, PJU, NN, MMD, JPA, EHM

Writing – review & editing: SP, EHM

## Data, code, and materials availability

All data are available in the main text or the supplementary materials.

## Supplementary Materials

**Figs S1-S14**

**Fig S1**: Gating hierarchy to identify IGRP-reactive CD8^+^ T cells in NOD mice.

**Fig S2**: Bulk RNA sequencing of IGRP-reactive CD8^+^ T cells from NOD mice.

**Fig S3**: Evaluation of anti-CD3 F(ab)_2_ for diabetes prevention in NOD mice.

**Fig S4**: Effect of anti-CD38 mAb on B cells in NOD mice.

**Fig S5**: Lymphodepletion analysis following anti-CD38 mAb and anti-CD3 F(ab)_2_ treatment in NOD mice.

**Fig S6**: Effect of anti-CD38 mAb and anti-CD3 F(ab)_2_ on memory T cells in NOD mice.

**Fig S7**: Evaluation of long-term effect of anti-CD38 mAb treatment in NOD mice.

**Fig S8**: Evaluation of CD38 expression on NOD pancreatic islets after anti-CD38 mAb treatment.

**Fig S9**: Co-expression of CD38 and SA-β-Gal in islets.

**Fig S10**: Refolding, and validation of HLA-A*02:01-restricted GAD65 and IGRP epitopes for spheromer assembly.

**Fig S11**: CD38 expression in IGRP-reactive CD8^+^ T cells in T1D patient samples.

**Fig S12**: Bulk RNA sequencing of CD38^+^ autoreactive T cells from recent-onset T1D patients.

**Fig S13**: Non-fasting blood glucose (NBG) profiles of NSH-HLA-A2/HHD mice.

**Fig S14.** Expression of CD38 in T cells in graft *versus* host disease (GVHD) patient samples.

