## Supplementary data for "CD38 defines a therapeutically targetable pathogenic T cell population for precision immunotherapy in autoimmune diabetes"

Everett H. Meyer, MD, PhD

Associate Professor,

Division of Blood and Marrow Transplantation and Cellular Therapy

Stanford University School of Medicine; Stanford, CA 94305, USA

### Supplementary Figures

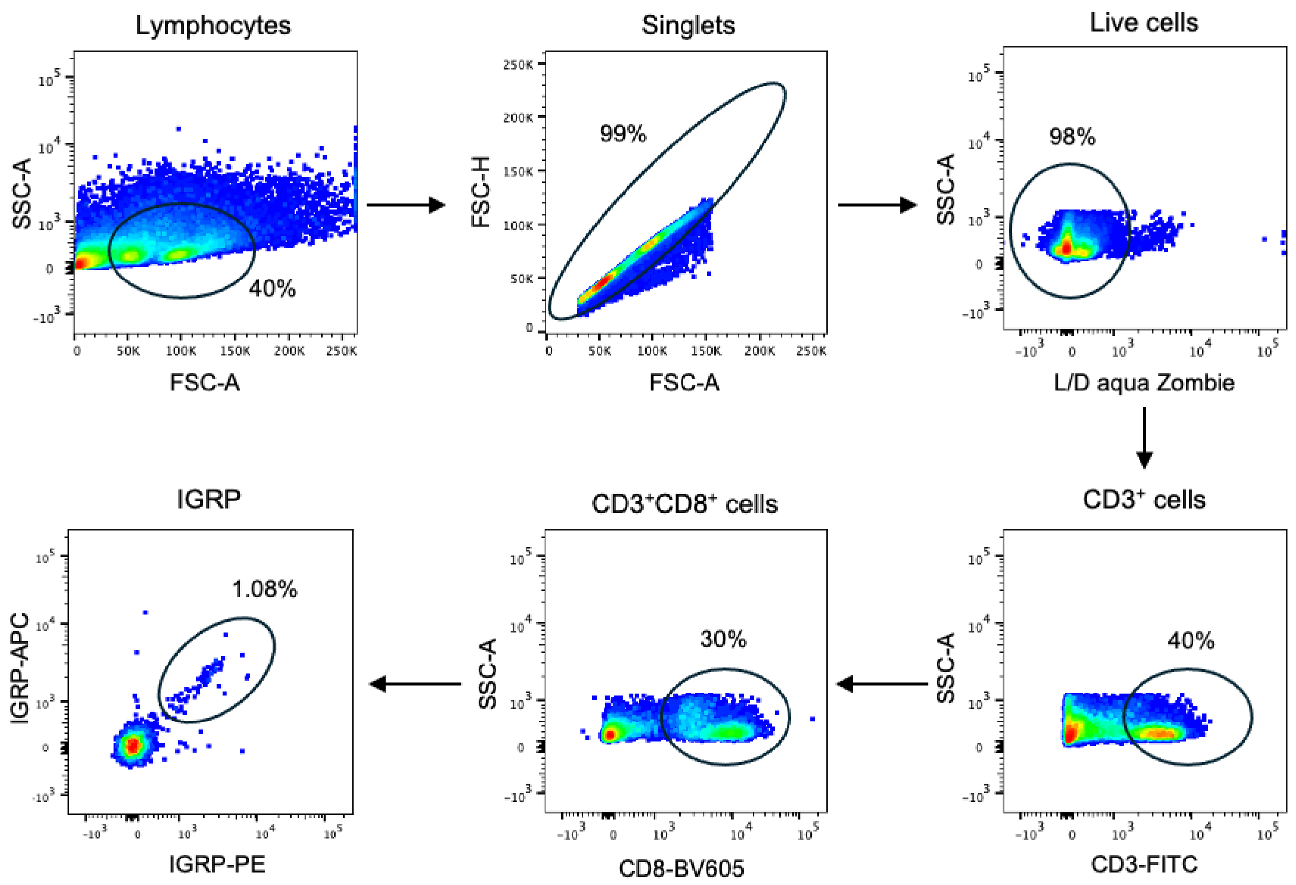


**Fig S1. Gating hierarchy to identify IGRP-reactive CD8⁺ T cells in NOD mice.** Lymphocytes were first gated by forward/side scatter, then Live/Dead Aqua-negative cells were selective as viable. From the viable cells, $CD3^+$ T cells were gated and subsequently restricted to the $CD8^+$ subset. Within $CD8^+$ T cells, IGRP-reactive cells were identified using APC- and PE-conjugated IGRP tetramers and gated as tetramer-positive events (dual-color gates).

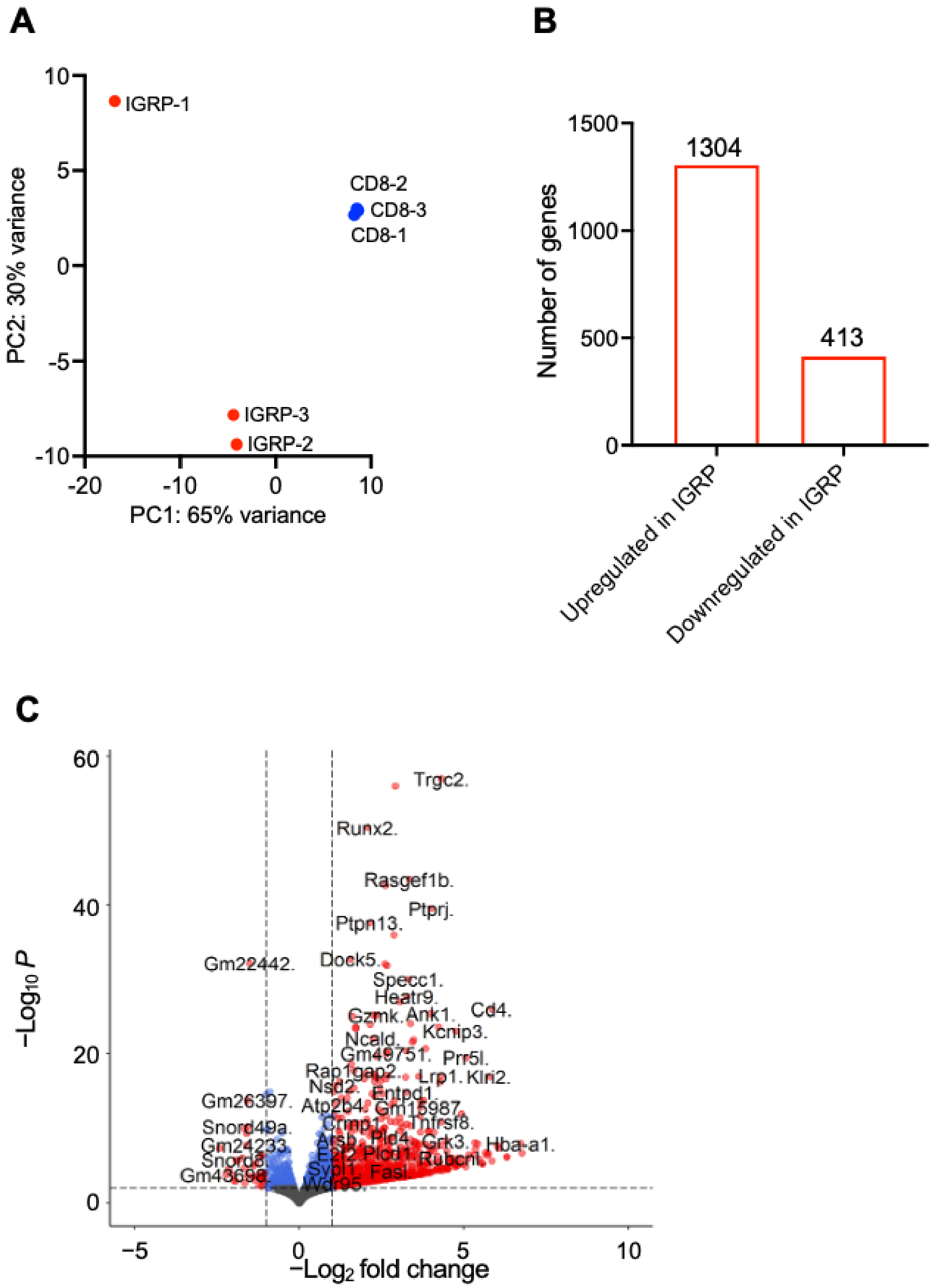


**Fig S2. Bulk RNA sequencing of IGRP-reactive CD8⁺ T cells from NOD mice.** (**A**) Principal component (PC) analysis of gene expression from IGRP-reactive $CD8^+$ T cells and bulk $CD8^+$ T cells. (**B**) Total number of upregulated and downregulated genes in IGRP-reactive $CD8^+$ T cells *vs* bulk $CD8^+$ T cells. (**C**) Volcano plot showing gene expression $\log_2$ fold change and –log10($P_{adj}$) in IGRP-reactive $CD8^+$ T cells *vs* bulk $CD8^+$ T cells.

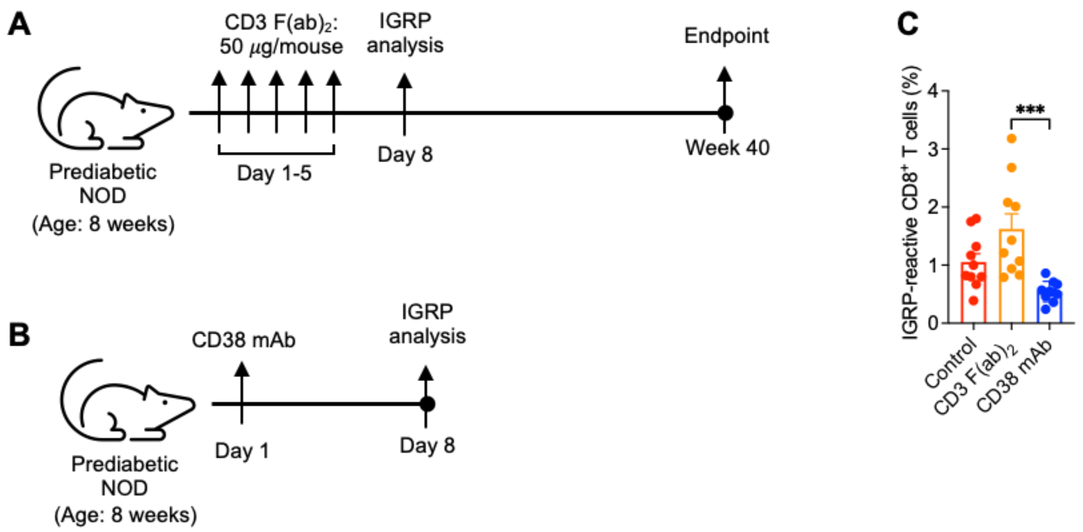


**Fig S3. Evaluation of anti-CD3 F(ab)2 for diabetes prevention in NOD mice.** (**A**) Schematic of the experimental design showing anti-CD3 F(ab)2 dosing in prediabetic female NOD mice: mice received intravenous 50 $\mu$g of anti-CD3 F(ab)2 for 5 consecutive days. Non-fasting blood glucose (NBG) was monitored weekly with a portable glucometer and followed through 40 weeks of age. (**B**) Schematic of the experimental design showing anti-CD38 mAb dosing in prediabetic female NOD mice: mice received single dose of 20 mg/kg of anti-CD38 mAb. (**B**) Percentage of IGRP-reactive $CD8^+$ T cells on day 8. Data were pooled from two independent experiments and represent mean ± SEM. ****p*<0.001.

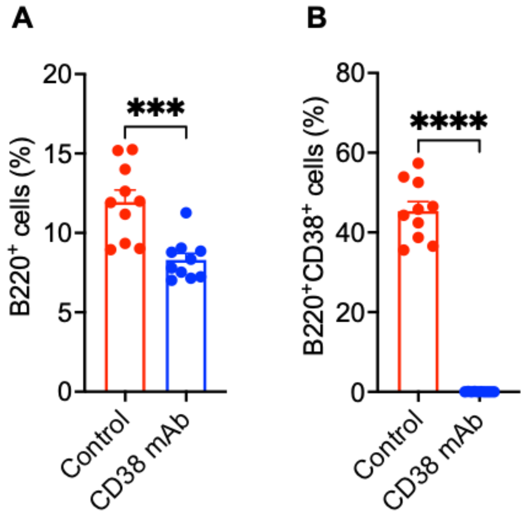


**Fig S4. Effect of anti-CD38 mAb on B cells in NOD mice.** Peripheral blood was collected from control and CD38 mAb treated mice at week 20 of age and stained with B220 and CD38 antibodies. (**A**) Percentage of B220$^+$ B cells. (**B**) Percentage of CD38$^+$ B cells. Data were pooled from three independent experiments and represent mean ± SEM. ***$p$<0.001, ****$p$<0.0001.

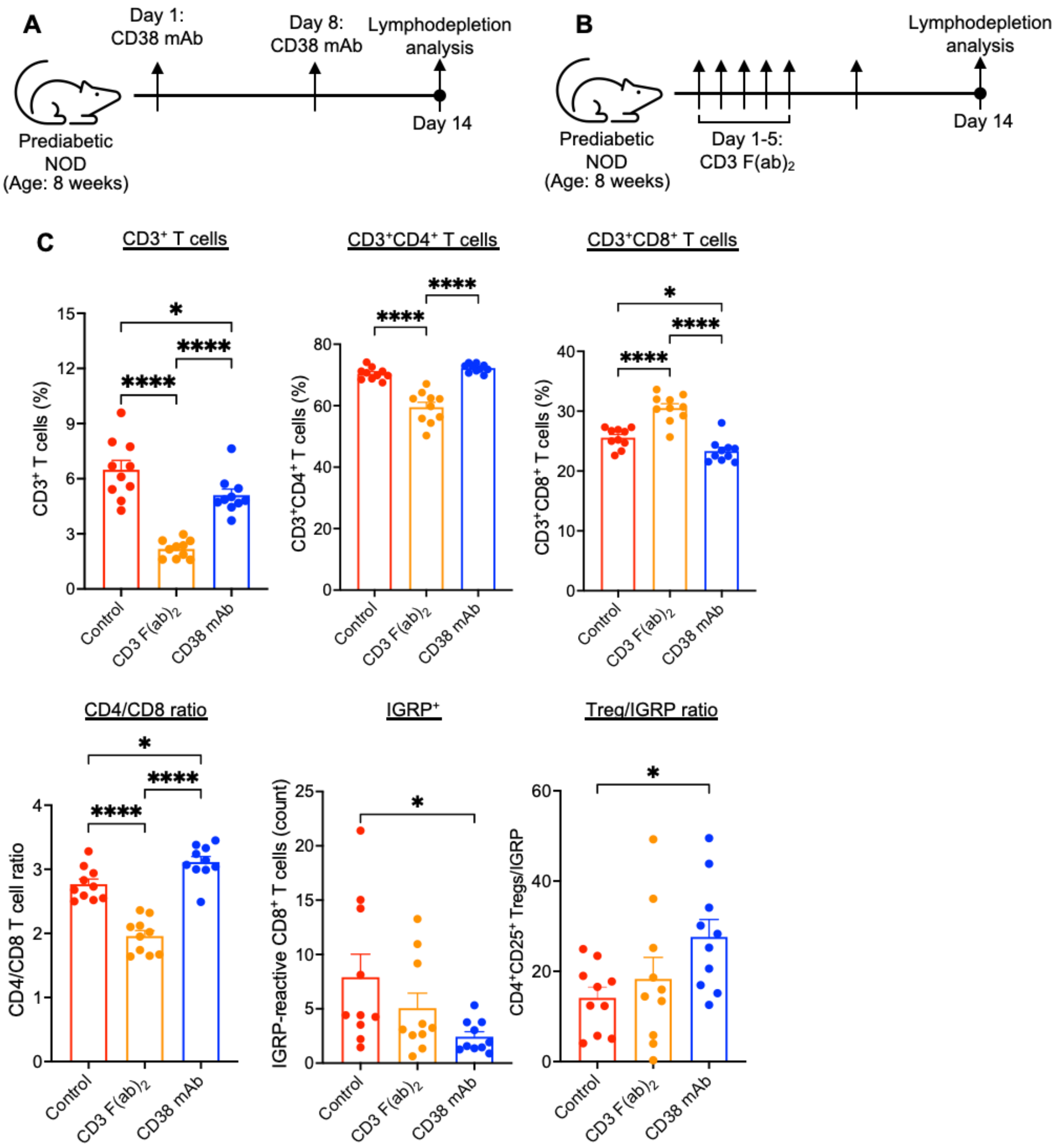


**Fig S5. Lymphodepletion analysis following anti-CD38 mAb and anti-CD3 F(ab)2 treatment in NOD mice.** Eight weeks old female prediabetic NOD mice were treated with two regimens. (**A**) anti-CD38 mAb was administered intraperitoneally at 20 mg/kg on days 1 and 8. (**B**) Anti-CD3 F(ab)2 was administered intravenously at 50 $\mu$g per mouse daily for five consecutive days (days 1-5). Peripheral blood was collected on day 14 and analyzed by flow cytometry using antibodies against CD3, CD4, CD8, CD25, and IGRP-tetramers to quantify circulating T cells and IGRP-reactive $CD8^+$ T cells. (**C**) Analysis of circulating T-cell subsets. Data were pooled from two independent experiments and represent mean ± SEM. **p*<0.05, *****p*<0.0001.

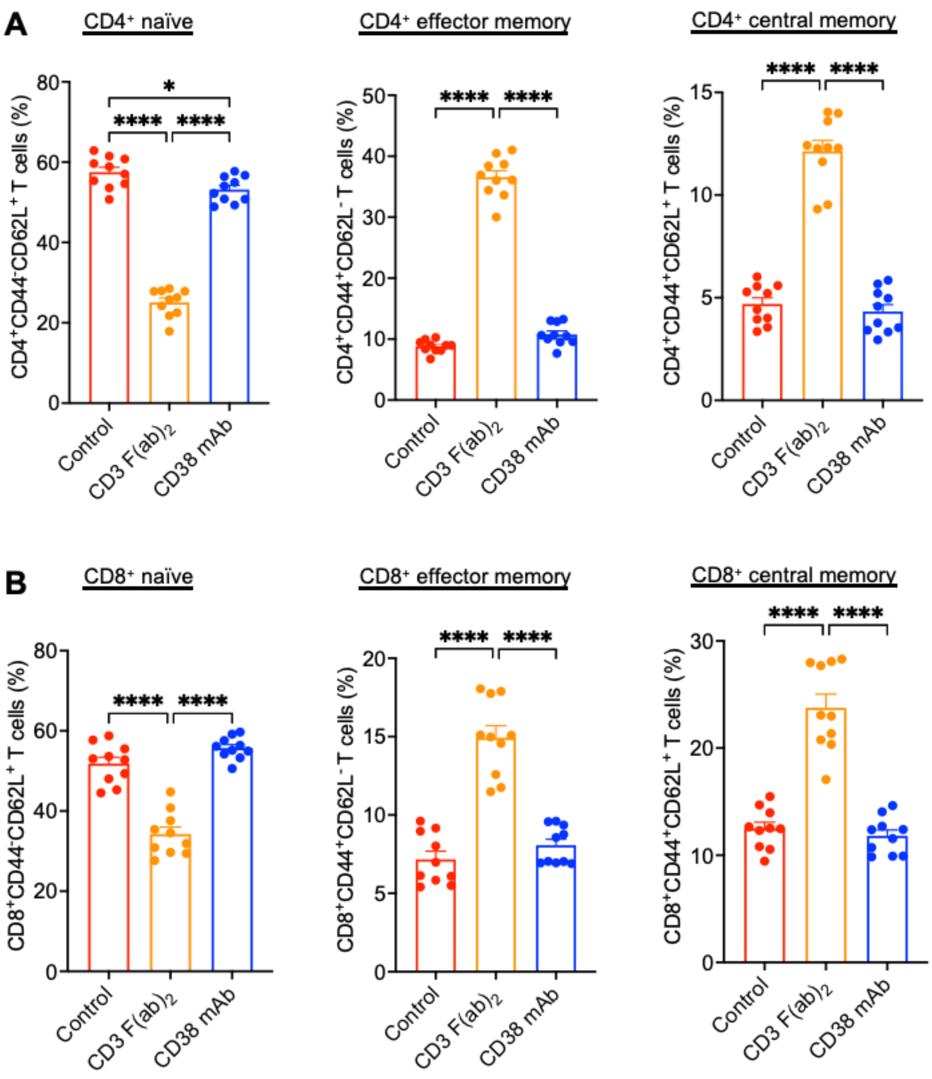


**Fig S6. Effect of anti-CD38 mAb and anti-CD3 F(ab)2 on memory T cells in NOD mice.** Eight weeks old female prediabetic NOD mice were treated with to regimens: anti-CD38 mAb given intraperitoneal at 20 mg/kg on days 1 and 8, or anti-CD3 F(ab)2 given intravenously at 50 $\mu$g per mouse daily for five consecutive days (days 1-5). Peripheral blood was collected on day 14 and analyzed by flow cytometry using antibodies against CD3, CD4, CD8, CD44 and CD62L. (**A**) Percentage of $CD4^+$ naïve ($CD44^-CD62L^+$), effector memory ($CD44^+CD62L^-$), and central memory ($CD44^+CD62L^+$) T cells. (**B**) Percentage of $CD8^+$ naïve ($CD44^-CD62L^+$), effector memory ($CD44^+CD62L^-$), and central memory ($CD44^+CD62L^+$) T cells. Data were pooled from two independent experiments and represent mean ± SEM. *$p$<0.05, ****$p$<0.0001.

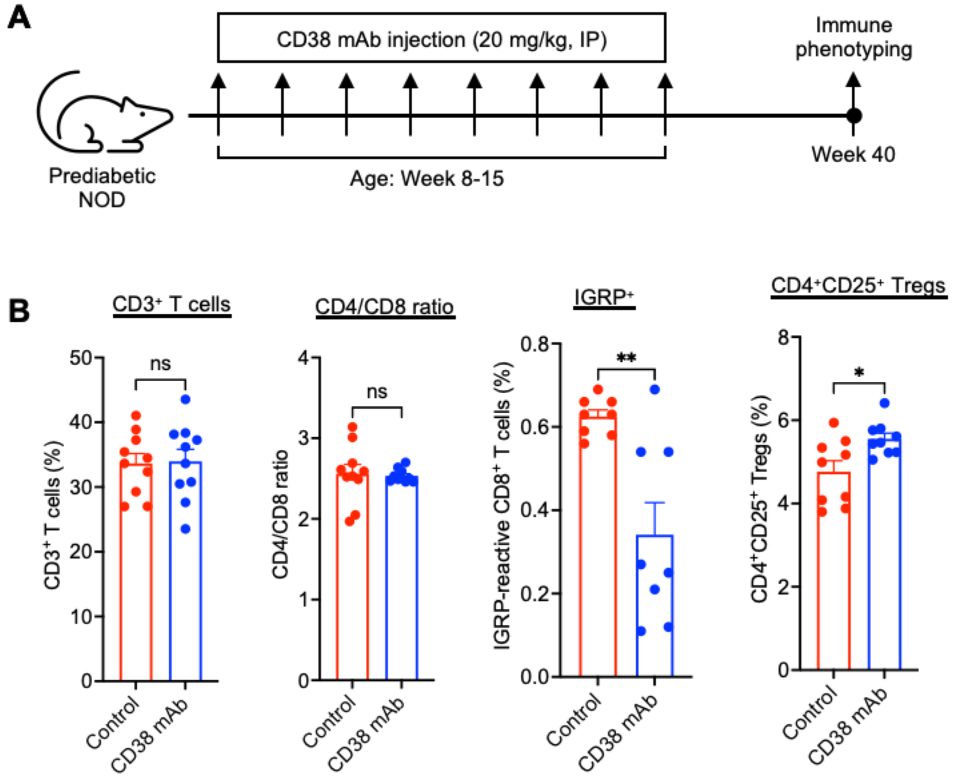


**Fig S7. Evaluation of long-term effect of anti-CD38 mAb treatment in NOD mice.** (**A**) Experiment design and timeline. Prediabetic NOD mice received eight intraperitoneal injections of anti-CD38 mAb once weekly from week 8-15. Mice were euthanized at week 40 for immunophenotyping. Vehicle treated NOD mice that survived to week 20 were used as controls. (**B**) Analysis of T cell subsets in spleen by flow cytometry. Data were pooled from two independent experiments and represent mean ± SEM. ns: not significant, *$p<0.05$, **$p<0.01$.

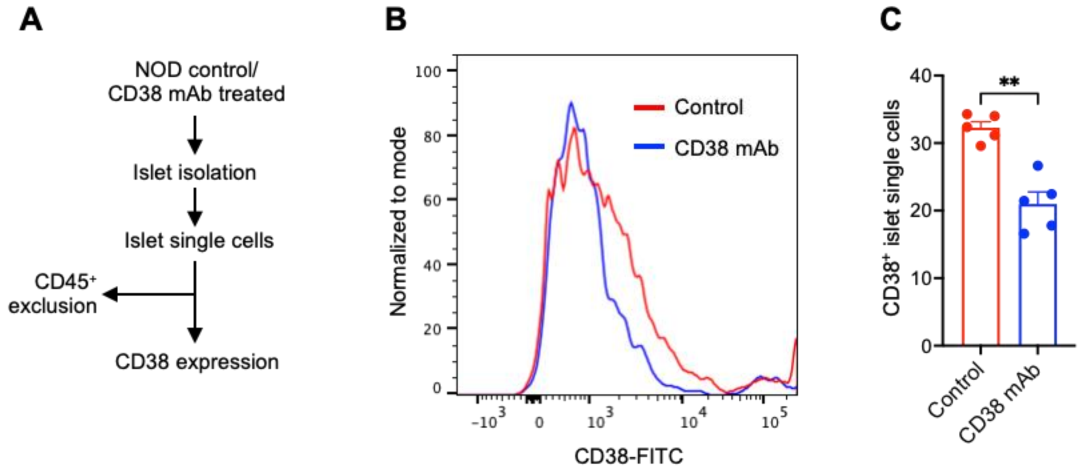


**Fig S8. Evaluation of CD38 expression on NOD pancreatic islets after anti-CD38 mAb treatment.** (**A**) Pancreatic islets were isolated from anti-CD38 mAb and vehicle treated NOD mice at week 20 of age. Purified islets were dissociated into single cells and stained with antibodies against CD45 and CD38. The expression of CD38 was analyzed among the $CD45^-$ live cells using flow cytometry. Data were pooled from two independent experiments and represent mean ± SEM. ***p*<0.01.

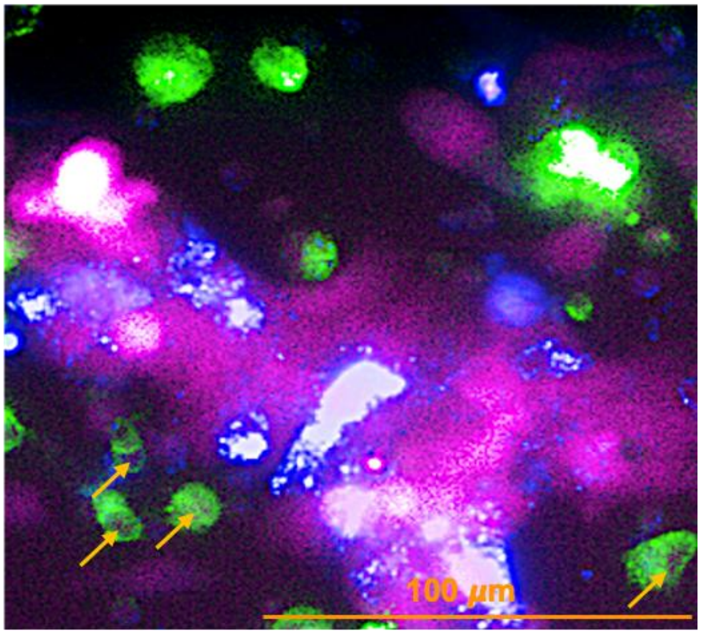


**Fig S9**. Co-expression of CD38 and SA-β-Gal in islets. Arrows indicate co-staining of CD38 and SA-β-Gal. (Image enlarged from **Fig 4I**). Magnification: 40×, Scale bar: 100 $\mu$m.

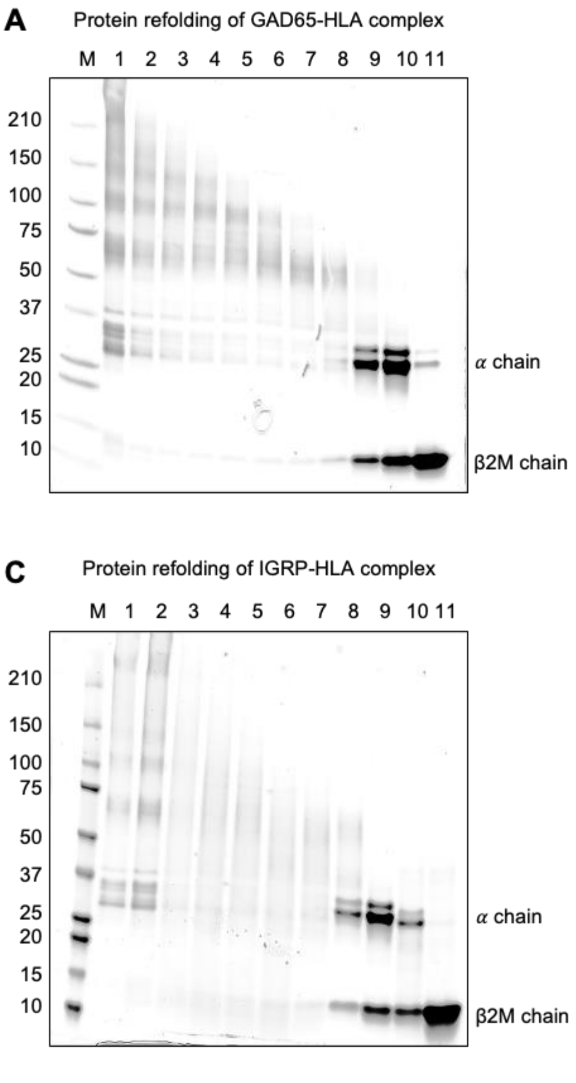


**Fig S10. Refolding, and validation of HLA-A*02:01-restricted GAD65 and IGRP epitopes for spheromer assembly.** Peptide epitopes were screened for predicted HLA-A*02:01 binding affinity using the IEDB analysis platform. GAD65 and IGRP formed stable pHLA complexes and were assembled into spheromer reagents. Refolded pHLA complexes were purified by size-exclusion chromatography (SEC), and fractions corresponding to properly folded monomeric pHLA complexes were confirmed by SDS-PAGE analysis. (**A**). Representative SDS-PAGE gel showing GAD65-HLA-A*02:01 refolding and purification. (**B**) Streptavidin gel-shift assay of GAD65 spheromer. (**C**) Representative SDS-PAGE gel showing IGRP-HLA-A*02:01 refolding and purification. (**D**) Streptavidin gel-shift assay of IGRP spheromer.

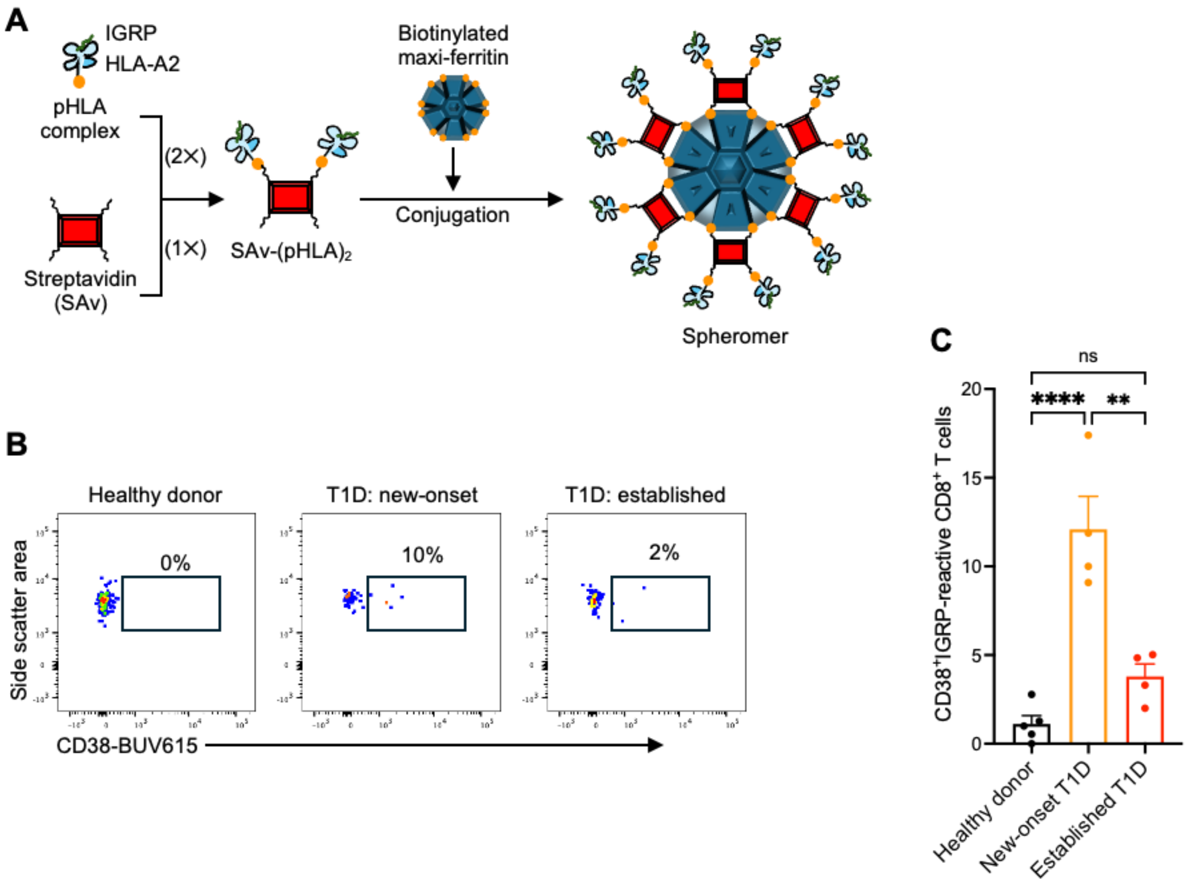


**Fig S11. CD38 expression in IGRP-reactive CD8⁺ T cells in T1D patient samples. (A)** Assembly workflow of HLA-A2 restricted IGRP-specific spheromer reagents. Biotinylated peptide-HLA-A02:01 (pHLA) monomers containing IGRP-derived epitopes were complexed with streptavidin (SAv) to generate intermediate SAv-$(pHLA)_2$ complexes. These intermediates were subsequently conjugated to a biotinylated 24-mer maxi-ferritin nanoparticle scaffold to assemble multivalent spheromer complexes. (**B**) Representative contour plots showing surface expression of CD38 among IGRP-reactive $CD8^+$ T cells. (**C**) Quantification of CD38-expressing IGRP-reactive $CD8^+$ T cells. Data represent mean ± SEM. ns: not significant, **$p$<0.01, ****$p$<0.0001.

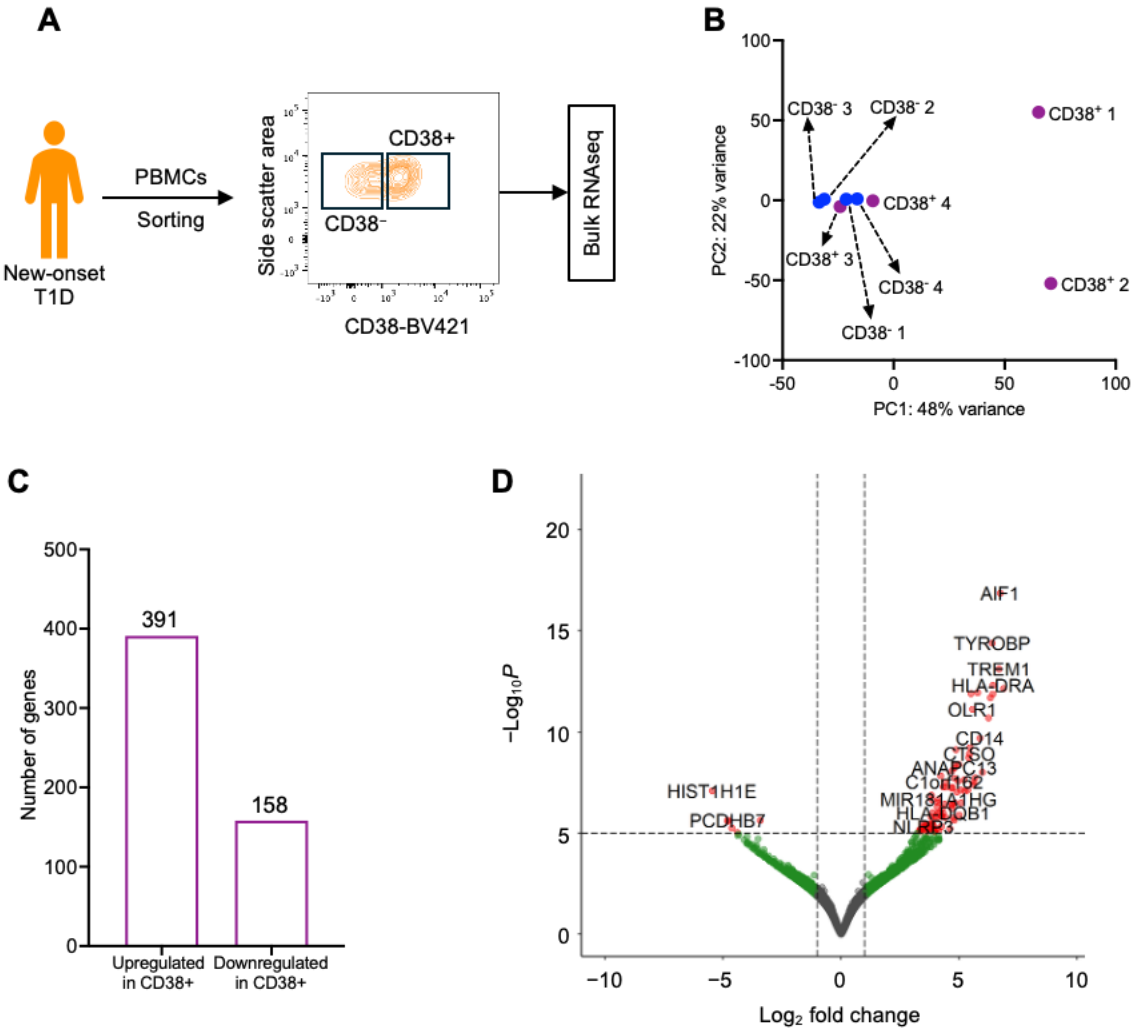


**Fig S12. Bulk RNA sequencing of CD38$^+$ autoreactive T cells from recent-onset T1D patients.** (**A**) Sorting of CD38$^+$ and CD38$^-$ autoreactive T cells from new-onset T1D patients. (**B**) Principal component (PC) analysis plot. (**C**) Number of upregulated and downregulated genes in CD38$^+$ autoreactive T cells *versus* CD38$^-$. (**D**) Volcano plot showing upregulated/downregulated genes in CD38$^+$ *versus* CD38$^-$ autoreactive T cells.

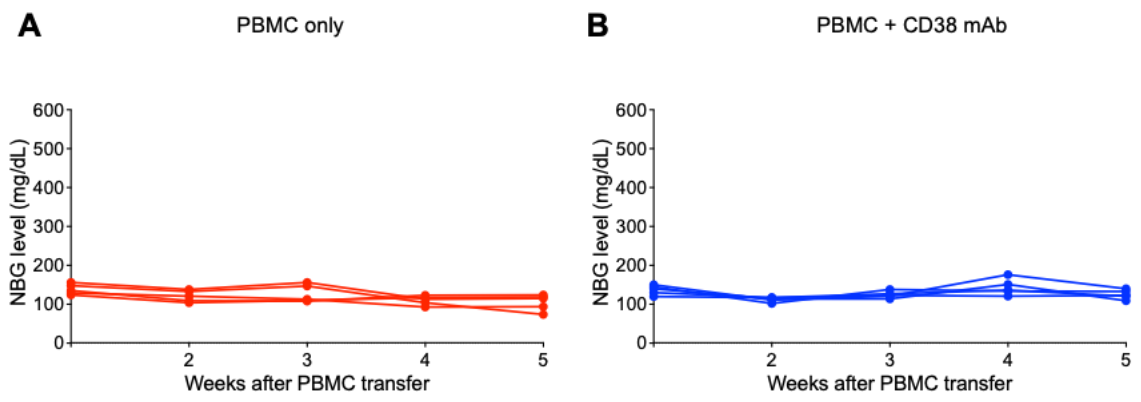


**Fig S13. Non-fasting blood glucose (NBG) profiles of NSH-HLA-A2/HHD mice.** (**A**) PBMC-only group. (**B**) PBMC + anti-CD38 mAb group.

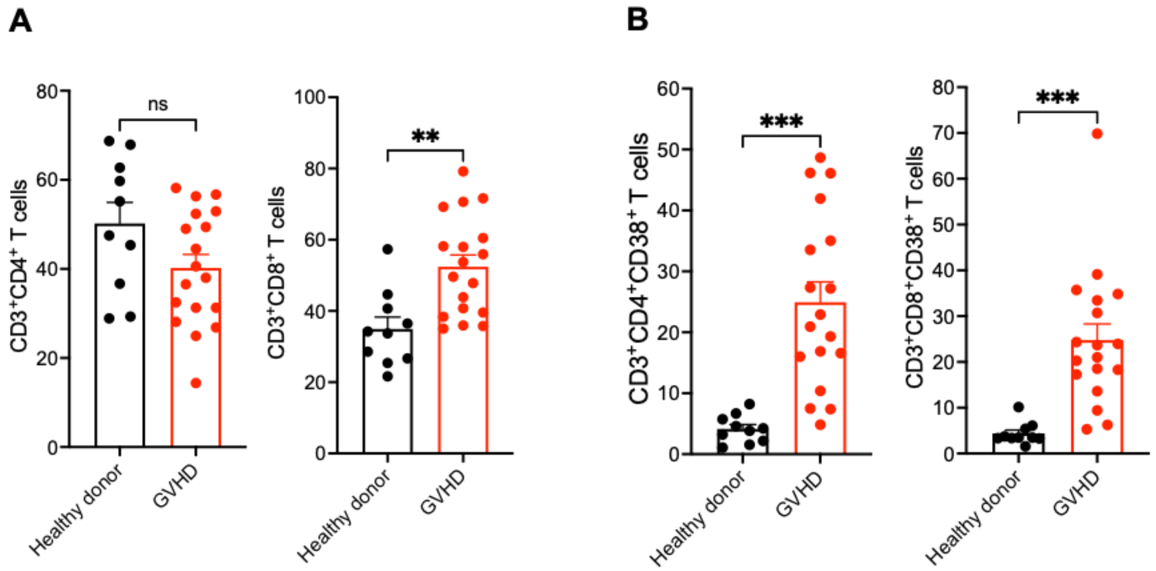


**Fig S14. Expression of CD38 in T cells in graft *versus* host disease (GVHD) patient samples.** Individuals with GVHD provided informed consent in accordance with the Declaration of Helsinki and were enrolled in a research protocol approved by the Stanford University Institutional Review Board (IRB) protocol #8903. (**A**) Frequency of $CD3^{+}CD4^{+}$ and $CD3^{+}CD8^{+}$ T cells in healthy donors and GVHD patients. (**B**) Percentage of $CD38^{+}$ T cells among $CD3^{+}CD4^{+}$ and $CD3^{+}CD8^{+}$ T cells in healthy donor and GVHD patients. Data represent mean ± SEM. ns: not significant, $^{**}p<0.01$, $^{***}p<0.001$.
